# Light and temperature regulate m^6^A-RNA modification to regulate growth in plants

**DOI:** 10.1101/2023.01.17.524395

**Authors:** Oliver Artz, Amanda Ackermann, Laura Taylor, Peter K. Koo, Ullas V. Pedmale

## Abstract

N6-methyladenosine is a highly dynamic, abundant mRNA modification which is an excellent potential mechanism for fine tuning gene expression. Plants adapt to their surrounding light and temperature environment using complex gene regulatory networks. The role of m^6^A in controlling gene expression in response to variable environmental conditions has so far been unexplored. Here, we map the transcriptome-wide m^6^A landscape under various light and temperature environments. Identified m^6^A-modifications show a highly specific spatial distribution along transcripts with enrichment occurring in 5’UTR regions and around transcriptional end sites. We show that the position of m^6^A modifications on transcripts might influence cellular transcript localization and the presence of m^6^A-modifications is associated with alternative polyadenylation, a process which results in multiple RNA isoforms with varying 3’UTR lengths. RNA with m^6^A-modifications exhibit a higher preference for shorter 3’UTRs. These shorter 3’UTR regions might directly influence transcript abundance and localization by including or excluding cis-regulatory elements. We propose that environmental stimuli might change the m^6^A landscape of plants as one possible way of fine tuning gene regulation through alternative polyadenylation and transcript localization.

## INTRODUCTION

Precise regulation of gene expression to specify growth, development and reproduction is paramount for the survival of all organisms. Therefore, complex regulatory mechanisms have evolved to precisely regulate gene expression through the entire life cycle of an organism. Methylation of DNA (Jones, 2012) and histones (Shi, 2007) are well understood regulators of gene expression. RNA can also be reversibly methylated on adenosine, known as N6-methyladenosine (m^6^A) (Jia et al., 2011; Zheng et al., 2013). m^6^A is the most abundant RNA modification (Dominissini et al., 2012), occurring in all major evolutionary lineages (Meyer and Jaffrey, 2017). However, the functions of the m^6^A mark and the underlying molecular mechanisms governing the regulation of m^6^A-modifications remain elusive. Transcript m^6^A-modification greatly influences diverse aspects of RNA metabolism such as RNA structure, maturation, processing, export, and translation (Zhao et al., 2016). Thus, m^6^A is an excellent candidate for regulating adaptive developmental responses through regulating gene expression. In plants, light and temperature are important developmental and adaptive signals which orchestrate processes such as seed germination, seedling de-etiolation, phototropic growth, shade avoidance, and floral transition (Kami et al., 2010). Plants grown in complete darkness undergo a developmental program called skotomorphogenesis or etiolation. Under these conditions, plants develop an elongated hypocotyl, apical hook, and closed, pale cotyledons. Upon light exposure, plants transition to photomorphogenesis or de-etiolation whereby hypocotyl elongation is inhibited, the apical hook opens, greening occurs and the cotyledons expand.

Plants perceive quality, quantity, duration, and periodicity of light through five distinct classes of photoreceptors: Cryptochromes (CRY1 and 2), Phytochromes (PHYA-E), Phototropins (PHOT1 and 2), UV-B RESISTANCE LOCUS 8 (UVR 8), and members of the ZEITLUPE (ZTL/LKP1/FKF1) family (Galvao and Fankhauser, 2015). A*rabidopsis CRY1 and CRY2* mediate responses to blue and UV-A light (Ahmad and Cashmore, 1993; Lin et al., 1996). Thus, without functional CRYs, plants are not able to effectively elicit phenotypic adaptations to blue light irradiation. While wild type (WT) seedlings grown in monochromatic blue light develop short hypocotyls and open green cotyledons, *cry1cry2* mutant seedlings develop elongated hypocotyls, reminiscent of dark-grown seedlings (Mockler et al., 1999).

The gene family of PHYs consists of five members (PHYA-E) which perceive light in the red to far red region (Sharrock and Quail, 1989). Within the family, PHYB is the most abundant member (Sharrock and Clack, 2002) with seedlings lacking functional PHYB exhibiting elongated hypocotyls when grown in monochromatic red light (Reed et al., 1993). A low red to far red-light ratio is created during shade conditions experienced under a vegetative canopy. Thus, *phy* mutants are also defective in the shade avoidance response (elongated hypocotyls, stems and petioles) (Ballare and Pierik, 2017). Phenotypic adaptions to shade closely mimic the adaptions to high temperatures (Delker et al., 2017) and the phenotypic responses to temperature and light share common molecular components including PHYB, which is known to function as a thermosensor, PIF4 transcription factor and the auxin biosynthetic genes SAV3/TAA1 and YUCCA (Franklin et al., 2011; Jung et al., 2016; Legris et al., 2016; Sun et al., 2012).

CRYs and PHYs regulate gene expression via protein-protein interactions, thus influencing the activity of transcription factors directly, or indirectly through enhancing transcription factor degradation (Galvao and Fankhauser, 2015). In metazoans, CRYs play a role in controlling m^6^A-modifications: Mice without functional CRYs exhibit lower levels of m^6^A-modified RNA and lose circadian regulation of m^6^A RNA abundance in liver samples (Wang et al., 2015). Moreover, in *Drosophila* m^6^A influences neuronal functions and sex determination (Lence et al., 2016).

mRNA is regulated through various mechanisms including expression, localization, splicing, and translation. Hence, mRNA function is not only dependent on abundance, but also localization of transcripts. Our current understanding of regulated mRNA localization has been gained from studies conducted in metazoans or microorganisms (Chin and Lécuyer, 2017; Holt and Bullock, 2009; Lazzaretti and Bono, 2017). For example, asymmetric distribution of transcripts is essential for the development of *Drosophila* embryos (Lécuyer et al., 2007) or *Xenopus* oocytes (King et al., 2005). In yeast, mRNA is transported from the mother to the daughter cell (Gonsalvez et al., 2005). Regulated mRNA localization correlates with the presence of cis-elements, so called “zip codes”, primarily in 3’UTRs (Hervé et al., 2004). In mammalian cells YTHDF2 (YTH DOMAIN FAMILY 2) binds specifically to m^6^A-modified transcripts and promotes their localization to processing bodies (P bodies) (Wang et al., 2014). Recently, it has been proposed that m^6^A-modifications might regulate RNA localization in the brain via controlling the interaction with specific RNA-binding proteins (RBP) (Madugalle et al., 2020).

To date our understanding of RNA localization in plants is limited: rice endosperms exhibit an uneven distribution of storage protein mRNAs on the endoplasmic reticulum (ER) (Li et al., 1993). Similar to other organisms, this localization is dependent on cis-elements mainly in 5’ and 3’UTRs (Washida et al., 2009). In addition to zip codes, the RBP OsTudor-SN controls mRNA localization in rice endosperm (Chou et al., 2017). Apart from being localized to the ER, plant mRNAs are spatially associated with mitochondria (Michaud et al., 2010, 2014a) and chloroplasts (Gómez and Pallás, 2010; Uniacke and Zerges, 2009). It has been reported that essential elements for those associations, particularly mitochondrial-associations, are localized in the UTRs of transcripts (Michaud et al., 2014b; Vincent et al., 2017). Moreover, intercellular RNA trafficking is essential for plant development and adaptive responses (Liu and Chen, 2018). Different types of small RNAs such as microRNAs (miRNAs), small interfering RNAs (siRNAs) or transfer RNAs (tRNAs) and mRNAs move directly between neighboring cells through plasmodesmata or systemically through the phloem (Chitwood et al., 2009; Kim et al., 2001; Zhang et al., 2009). Many mobile transcripts in *Arabidopsis* possess tRNA-like structures that mediate RNA mobility in grafting experiments. These structures are present in coding and UTR regions of transcripts (Zhang et al., 2016). The importance of UTRs for RNA trafficking is underlined by findings from potato. Here, UTRs of *BEL5*, a BEL1-like transcription factor, are essential for long-distance mobility of *BEL5* transcripts (Banerjee et al., 2009). Taken together, this emphasizes the importance of mRNA localization and the role of UTRs in this respect for appropriate gene function.

One important process to control the length of 3’UTR is alternative polyadenylation (APA), which has been identified as one of the mechanisms by which m^6^A regulates genes. APA is a process of mRNA processing in which different polyadenylation sites are being used for a given transcript, thus producing alternative isoforms with varying 3’UTR lengths. In humans and plants, approximately 70% of genes use multiple polyadenylation sites (Derti et al., 2012; Wu et al., 2011). The majority of APA sites were found to be in close proximity of the stop codon (Shen et al., 2008). Hence, APA may lead to exclusion or inclusion of essential regulatory elements such as microRNA (miRNA)-binding sites or zip codes (Rhoades et al., 2002) and thus directly influence mRNA translation and abundance. Indeed, APA is an important way for plants to regulate their growth and development in response to environmental stimuli, thus controlling crucial developmental processes such as floral transition (Hornyik et al., 2010; Liu et al., 2010; Simpson et al., 2003), oxidative stress responses (Zhang et al., 2008), maintenance of circadian rhythm (Liu et al., 2014), seed dormancy (Cyrek et al., 2016), and auxin-mediated growth responses (Hong et al., 2018).

In plants, m^6^A-modifications play an essential role in growth and development. Processes such as embryo (Bodi et al., 2012; Zhong et al., 2008), leaf (Arribas-Hernandez et al., 2018; Arribas-Hernández et al., 2020; Růžička et al., 2017; Shen et al., 2016), trichome (Arribas-Hernandez et al., 2018; Wei et al., 2018), and root (Chen et al., 2018; Růžička et al., 2017) development as well as floral transition (Duan et al., 2017; Song et al., 2021) are greatly influenced by m^6^A modifications. However, very little is known about environmental regulation of RNA modifications, in particular m^6^A. The dynamic nature of m^6^A-modifications makes it an excellent candidate to regulate gene expression in response to variable conditions and many of the growth and developmental responses listed above, which are known to be influenced by m^6^A marks, are also known to be regulated by light and temperature.

In this study, we provide evidence that environmental stimuli shape the m^6^A RNA landscape in the model plant *Arabidopsis thaliana*. Our data suggest that region-specific methylation of transcripts might be important for RNA and protein localization and that the presence of m^6^A in 3’UTR regions of transcripts might lead to a shift in APA resulting in a higher preference for shorter 3’UTR regions. We conclude that this shift in APA site preference might be a means to fine tune gene expression, thus adapting to environmental conditions. Moreover, our analysis of differentially methylated transcripts enabled us to identify potentially new genes involved in specific pathways. For example, we identified *TAF15b* (*TATA-BINDING PROTEIN-ASSOCIATED FACTOR 15b*) to promote apical hook formation in etiolated seedlings, which is a so far undescribed role for this gene.

## RESULTS

### Identification of differential m^6^A-modification during adaptive responses

To investigate dynamic m^6^A-modified RNA underlying light and temperature-induced transcriptomic changes in *Arabidopsis*, we catalogued m^6^A-modified mRNA from seedlings grown in experimental conditions where morphological changes are readily observed. We chose the shade avoidance syndrome (SAS), thermomorphogenesis/high-temperature (HT), and de-etiolation for this purpose. Readily visible and quantitative features of SAS and HT are the increase in hypocotyl length, whereas de-etiolating seedlings exhibit hypocotyl growth inhibition (Figures 1A and S1A-S1C) (Delker et al., 2017; Neff and Chory, 1998; Smith and Whitelam, 1997). We also included *Arabidopsis* mutants lacking PHYB, CRY1 and CRY2 photoreceptors as they are impaired in their blue or red light perception, respectively (Figures 1A and S1A-C) (Mockler et al., 1999; Reed et al., 1993). Hence, *cry1cry2* mutants grown in blue light and *phyB* mutants grown in red light display a partially etiolated phenotype with characteristic elongated hypocotyls (Figures S1B-S1C).

**Figure 1.**
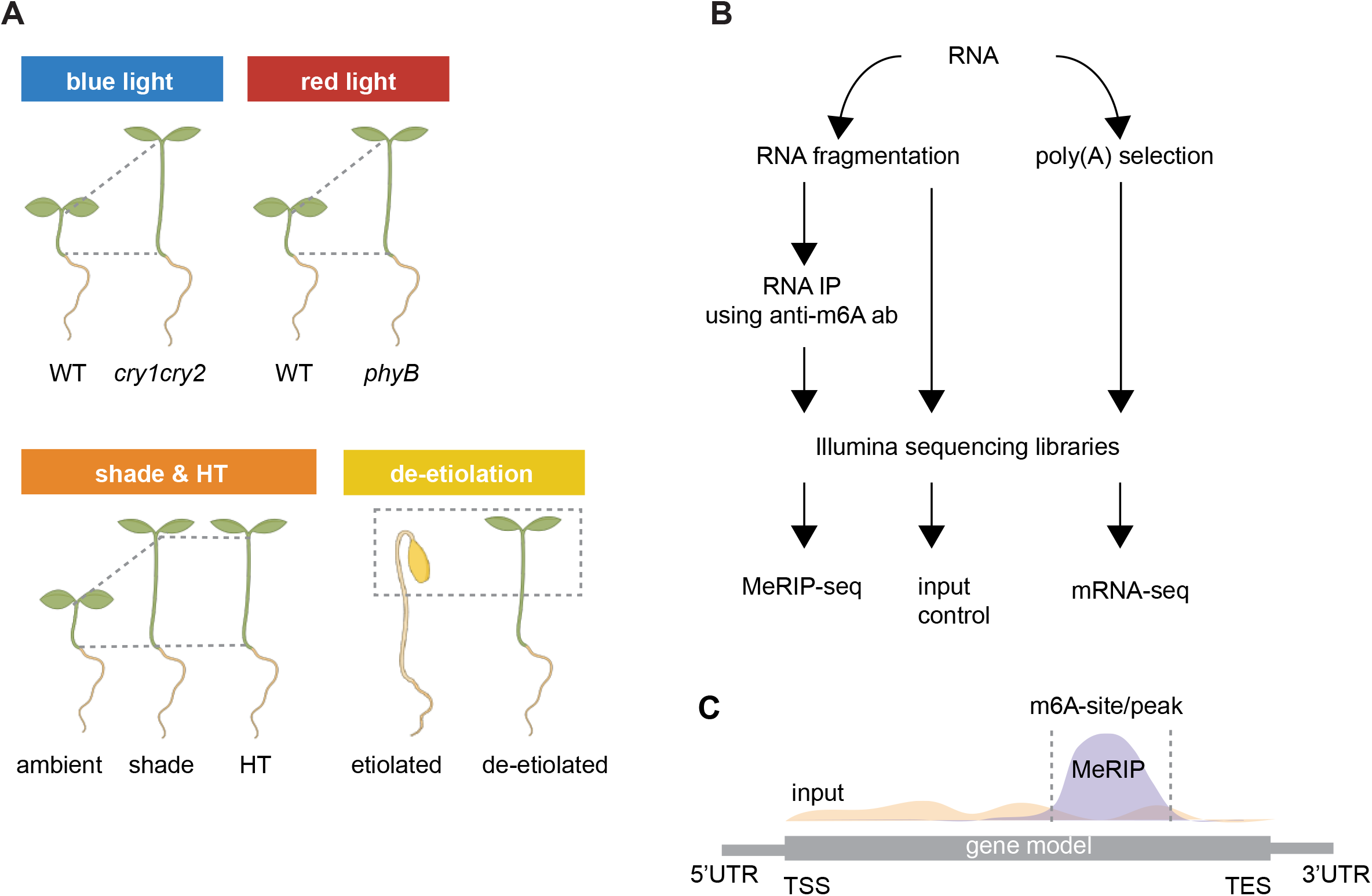
Experimental approach for identification of differential m^6^A RNA modification. **(A)** Schematic diagram illustrating plant growth conditions used in this study. WT and *cry1cry2* mutant plants were grown in constant blue light (10 μmol m^-2^ s^-1^) and WT and *phyB* mutant plants were grown under constant red light (20 μmol m^-2^ s^-1^) for 4 days. Another set of plants was grown in continuous white light at standard ambient temperature conditions (95 μmol m^-2^ s^-1^ white light, red:far-red ratio: 5.4, 22°C) and then shifted to either shade (red:far-red ratio: 0.2) or high temperature (HT) (30°C) conditions for 24 h before RNA extraction. For the de-etiolation assay, plants were grown in darkness for 4 days prior to a 24 h white light treatment (95 μmol m^- 2^ s^-1^) or additional 24 h in darkness. **(B)** Diagram illustrating the experimental pipeline. Total RNA was extracted from plant tissue and chemically fragmented into ∼200-nt-long fragments. RNA fragments were subjected to m^6^A-specific RNA immunoprecipitation with subsequent sequencing (MeRIP-seq). In addition, poly(A) selection was performed on the total RNA to isolate mRNA for sequencing (RNA-seq). **(C)** Schematic diagram illustrating identification of the m^6^A peak enrichment on the RNA when compared to its input control sample. Grey boxes represent the gene model with its 5’ and 3’UTR. TSS: Transcriptional start site; TES: Transcriptional end site.

Since m^6^A is biochemically similar to unmodified adenosine and is not easily detected by current sequencing techniques or bisulfite chemical modification, we used an anti-m^6^A antibody that specifically recognizes N-methyladenosine specifically to enrich for m6A-modified RNA (Bodi et al., 2010; Meyer et al., 2012). To this end, total RNA was extracted from seedlings treated as described in Figure 1A, was randomly fragmented to approx. 200-nt-long fragments and then enriched for m^6^A-modified RNAs by immunoprecipitation (MeRIP) (Dominissini et al., 2012; Meyer et al., 2012) (Figure 1B). The sequences of the immunoprecipitated RNA along with those of the input RNAs were determined using short-read RNA-sequencing (MeRIP-seq). Input RNA was used to normalize for the identification of enriched m^6^A-sites (Figure 1C) in the MeRIP-seq samples (Dominissini et al., 2012; Meyer et al., 2012). Along with MeRIP-seq, we also performed RNA-seq on poly(A)-purified RNA from the same total RNA source to gain more detailed information about gene regulation and to assess overall transcript abundance.

### m^6^A-modifications are dynamic in plants exposed to various adaptive growth conditions

The RNA-seq, input fragmented RNA, and MeRIP-seq sequencing reads were mapped to the *Arabidopsis thaliana* genome (version TAIR10). Our sequencing data from biological replicates were highly correlated, with a Pearson correlation coefficient of *r* ≥ 0.93 (Figure S2A). First, we examined the transcript levels of known marker genes for each experimental condition to confirm that the employed conditions elicited the expected transcriptional responses (Figure S2B). For example, *PHYTOCHROME INTERACTING FACTOR3-LIKE 1 (PIL1)* is a known marker gene for low red:far-red shade response and *YUCC8* (*YUC8*) for high temperature growth (Salter et al., 2003; Sun et al., 2012). We observed that *PIL1* and *YUC8* transcript levels increased by 20 and 2-fold in plants exposed to shade and HT, respectively, as expected (Figure S2B). Similarly, all treated seedlings exhibited the expected visible phenotypes (Figures S1B-C and S2B). These combined data indicate that our biological replicates were robust.

MeRIP-seq reads were compared to their input RNA sequencing reads using a peak calling software (exomePeak; (Meng et al., 2013)) with multiple replicates, where the enriched m^6^A-sites show peaks along the gene body (Figures 1C and 2A). We identified m^6^A-sites with a minimum false discovery rate (FDR) of <0.05. In our initial survey of m^6^A distribution within the transcriptome of our nine samples in five conditions, we identified a total of 23,435 m^6^A-sites in the RNAs of 5911 unique genes, representing approx. 20% of the known protein-coding genes in *Arabidopsis* (Figure 2B). Unbiased hierarchical clustering analysis on these unique 5911 genes identified distinct nodes that were grouped by experimental groups (Figure 2C). We found between 257 and 4978 m^6^A-sites for each tested condition (Figure 2C). A relatively low number of m^6^A-modified transcripts was observed in red light (424 sites in WT, 257 sites in *phyB* mutants) or darkness (etiolated, 920 sites) indicating that the specific light environment might have a large influence on the RNA methylome (Figure 2B). Blue light might be an important signal to induce m^6^A modification of transcripts since plants exposed to either monochromatic blue light or white light, which contains large quantities of blue light, show higher numbers of m^6^A-modified transcripts: 1977 sites were identified in de-etiolated plants which were exposed to white light and 4978 sites were identified in WT plants grown in blue light. In our dataset, m^6^A-modifications were identified on transcripts of genes that were lowly (FPKM < 3) or highly (FPKM > 1000) expressed (Figure S3A) indicating that our experimental setup is suitable to detect m^6^A-modified transcripts within a wide range of expression levels. We identified varied patterns of m^6^A-sites in all our samples, with a majority of genes containing a single site (Figure 2D), in agreement with a previous study (Luo et al., 2014). Highly differentially m^6^A-modified transcripts, which are exclusively m^6^A-modified in one of the experimental pairs were identified (Figure S3A). Interestingly, differentially m^6^A-modified transcripts can be significantly up-regulated or down-regulated between the sample pairs (Figure S3A). These results align with previous studies presenting evidence for the stabilizing and destabilizing effect of m^6^A on mRNA in plants (Anderson et al., 2018; Martinez-Perez et al., 2017; Parker et al., 2020; Shen et al., 2016; Wei et al., 2018). Our results suggest that m^6^A might regulate specific transcripts via different molecular pathways, in different tissues or different conditions.

**Figure 2.**
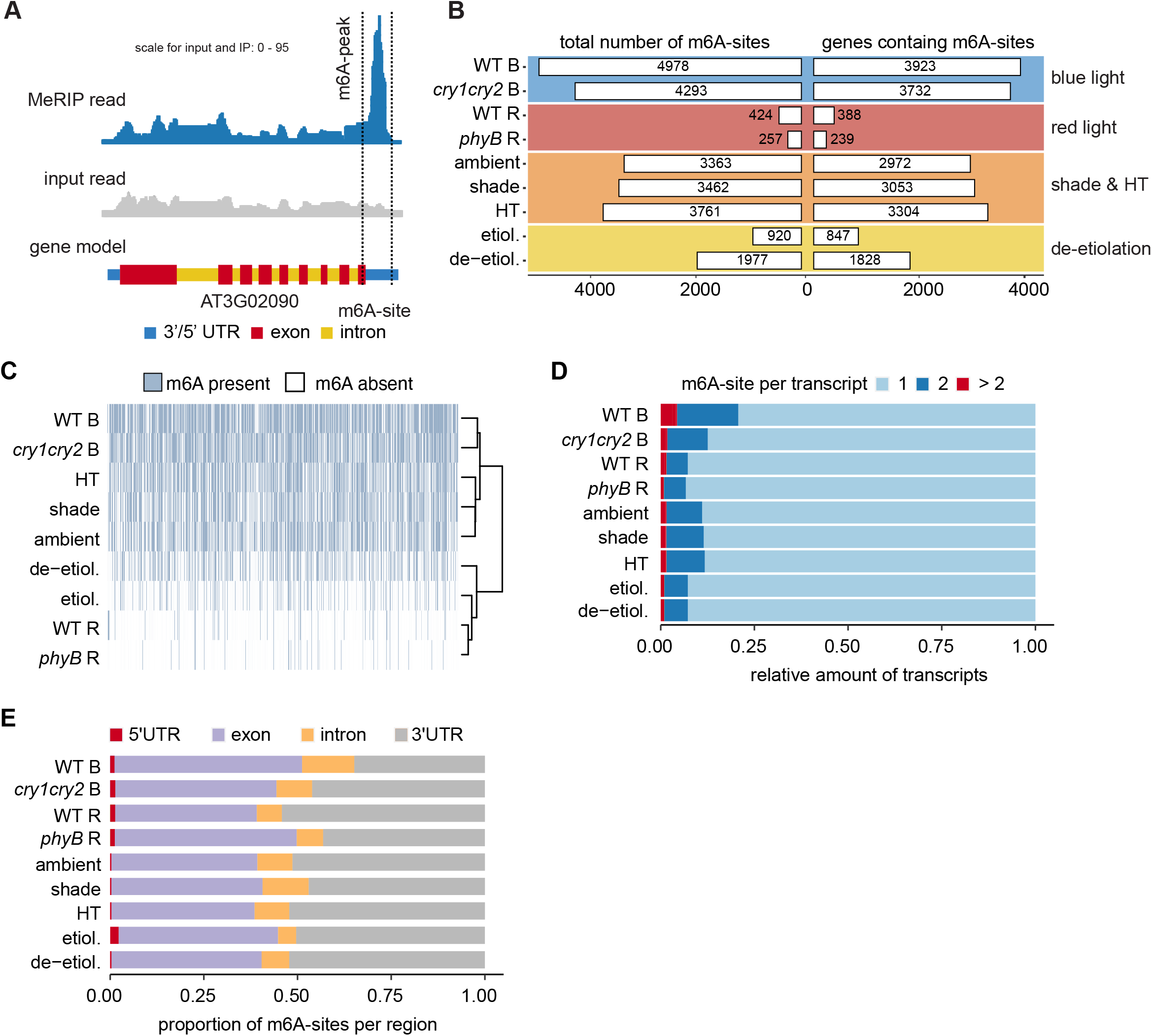
m^6^A-modifications are present in plants exposed to various environmental conditions. **(A)** Representative Arabidopsis gene plots harboring an m^6^A-site for AT3G02090. Normalized coverage of sequencing reads for input and MeRIP samples are shown in grey and blue. The blue peak within the dashed lines represents a highly significant m^6^A-site in the 3’UTR of the transcript. **(B)** The total number of identified m^6^A-sites for each condition is compared to the number of corresponding genes containing one or more m6A-sites in the indicated genotypes and experiments. **(C)** Dendrogram depicting the unique 5911 RNA of genes containing one or more m^6^A-sites across the genotypes and experiments. Samples cluster according to experimental pairs. **(D)** The number of m^6^A-sites identified per transcript are shown for the indicated conditions and genotypes. Most transcripts harbored a single m^6^A site. **(E)** Proportion of m^6^A-modifications identified within 5’UTR, exon, intron or 3’UTR regions of the transcripts in the indicated samples. The majority of m^6^A sites was found in exons or 3’UTRs. **(F)** Intersection of m^6^A-modified regions from all samples sorted by size. Circles indicate sets being part of the respective intersection. The set size describes the total number of identified modifications per region combined from all samples. Most transcripts harbor a single site either in 3’UTRs or exons. Very few transcripts harbor modifications in different regions.

We next examined the pattern of m^6^A-site distribution by categorizing each site as 5’UTR, exon, intron or 3’UTR of the respective transcript. We found that the majority (88%) of identified m^6^A peaks were found in 3’UTR and exon regions (Figures 2E). These combined findings suggest that m^6^A-modification varies and is dynamic across our experimental conditions and genotypes, possibly having a role in gene regulation during adaptive processes.

### m^6^A is highly enriched in the protein-coding and UTRs of transcripts

To further investigate if m^6^A modifications mainly act to control transcript abundance, we compared differentially expressed genes (DEG; FDR < 0.05) obtained from RNA-seq experiments with m^6^A-modified genes. We observed that on average less than 9% of DEGs and m^6^A-modified genes overlap (Figure 3A), which suggests that regulating transcript abundance might not be the most important way m^6^A influences gene expression. It is conceivable that specific m^6^A-modification of one or multiple sites on the mRNA have different effects on gene regulation. Hence, we investigated whether the used genotypes or environmental conditions have an effect on the overall pattern of m^6^A modification. To visualize enrichment patterns of m^6^A-sites at functionally important regions of genes, we performed metagene analyses. m^6^A was strongly enriched close to the transcriptional end site (TES) and less strongly at the transcriptional start sites (TSS) in all our samples (Figure. 3B). We did not observe a notable effect of the genotype or environment on the distribution of m^6^A-modification along transcripts by genotype or environmental conditions.

**Figure 3.**
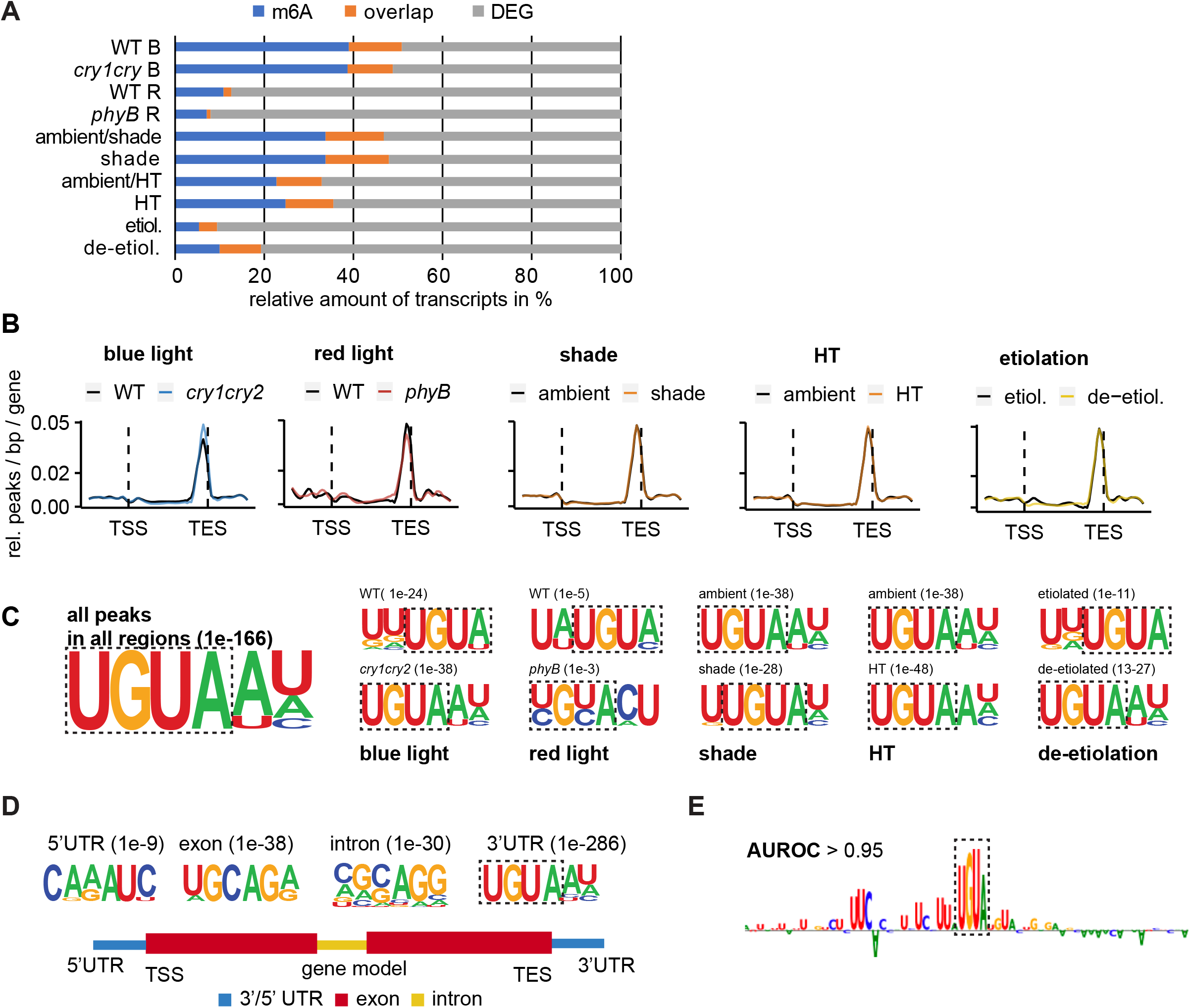
m^6^A is highly enriched around the TES within the UGUA motif. **(A)** Overlap of differentially expressed genes and m^6^A-modified genes are shown for each sample. A relatively small fraction of transcripts is differentially methylated and differentially expressed. **(B)** Metagene analysis showing MeRIP-seq read enrichment along transcripts. Sequences 5000 bp upstream of the transcription start site (TSS) and 5000 bp downstream of the transcription end site (TES) were considered in the analysis. m^6^A was highly enriched close to the TES in all condition and genotypes. **(C)** m^6^A-containing sites were analyzed for enriched RNA sequences using HOMER. The highest-ranking motifs are shown together with the corresponding p-value. m^6^A regions were highly enriched in the UGUA motif in all conditions and genotypes. **(D)** Enriched RNA sequences of m^6^A-containing sites were analyzed with respect to their location on transcripts. The highest-ranking motifs are shown together with the corresponding p-value. TSS: transcriptional start site, TES: transcriptional end site. The UGUA motif was significantly enriched in 3’UTR sites, but not other regions of transcripts. **(E)** Sequence logo of a representative attribution map generated by a deep learning model for a given mRNA sequence. Heights correspond to the importance of the nucleotide with a sign that corresponds to its impact on predictions. All peaks found in 3’UTRs were analyzed using a deep learning approach. Performance was measured by AUROC. The UGUA motif was significantly enriched in 3’UTR m^6^A regions.

Previous studies found the RRACH motif [(A/G) (A/G) AC (A/C/U)] to be enriched around m^6^A-sites in mammals (Dominissini et al., 2012; Meyer et al., 2012), insects, yeast (Schwartz et al., 2013), and plants (Anderson et al., 2018; Duan et al., 2017; Luo et al., 2014; Parker et al., 2020; Shen et al., 2016). However, additional motifs have been proposed to be important for m^6^A-modification: In Arabidopsis and maize, the URUAY motif [(U (A/G) UA (C/U)] was found to be enriched in addition to the RRACH motif (Miao et al., 2019). Moreover, the URUAY motif is the only enriched binding motif for the m^6^A reader protein ECT2 in Arabidopsis (Wei et al., 2018). We performed *de novo* motif identification and found that m^6^A was enriched in the context of a UGUA motif [UGUA (A/U) (U/A/C)] (Figure 3C, E, Figure S4). When we further investigated different transcript regions, we found that the UGUA motif was significantly enriched only in 3’UTR regions but not in 5’UTR, intron or exon regions of transcripts (Figure 3D). However, we found other motifs containing a dominant adenosine (A), which is likely to be methylated (5’UTR: C (A/G) (A/G) AU (C/U), exon: (U/A) GCAG (A/G), intron: (C/A/G/U) (G/A/C/U) (C/G/A/U) (A/C/U/G) (G/A/C) (G/C/A/U)). This suggests that there might be a region-specific motif for m^6^A-modification. We also used a deep learning approach to detect enriched RNA motifs, confirming enrichment of the UGUA motif in 3’UTR regions of m^6^A-modified transcripts (Figure 3E, Figure S6). Notably, while we found RRACH consensus sequences in our peaks, they were not identified to be significantly enriched in our dataset. This indicates that plants might have evolved a specific sequence context in which RNA is m^6^A-modified. Our results underline the notion that URUAY/UGUA is the most important motif for transcript m^6^A modification in plants, particularly in the 3’UTR of transcripts. However, other motifs might be important in other regions of transcripts.

### RNAs coding for chloroplast-targeted proteins show a distinct methylation pattern

m^6^A has multiple different functions and acts via different molecular mechanisms in plants (Arribas-Hernández and Brodersen, 2019). We further investigated the role of the position of m^6^A on transcripts. Position-specific functions have been described in mammalian cells where m^6^A-modifications in 5’UTRs promote cap-independent translation (Meyer et al., 2015). To analyze whether m^6^A-modification of different transcript regions correlates with specific biological processes, we grouped methylated transcripts from all samples and binned our data into different categories representing 5’UTR, exon, intron, and 3’UTR regions. Gene Ontology (GO) term enrichment analyses showed distinct and overlapping enriched GO terms between regions. Many general metabolic processes such as fatty acid metabolism, gene regulation or cellular nitrogen compound metabolic process were primarily enriched in both 3’ UTR and exons (Figure 4A). However, we identified a subgroup of GO terms, that were exclusively enriched in either exons or 3’UTRs: Among others, proteasomal protein catabolic process, Golgi vesicle transport, and vacuole organization were exclusively enriched in transcripts with m^6^A-modifications in 3’UTR regions. This suggests that the control of proteasomal protein turnover and the endomembrane system might mainly rely on the regulation of transcripts modified specifically in the 3’UTR or not-modified in exons. The GO terms auxin efflux, chlorophyll biosynthetic process, and chloroplast organization were specifically enriched in transcripts with m^6^A-modifications in exons, thus suggesting that auxin transport and chloroplast development might be regulated through transcripts with m^6^A-modifications in exons or lacking m^6^A-modifications in 3’UTRs.

**Figure 4.**
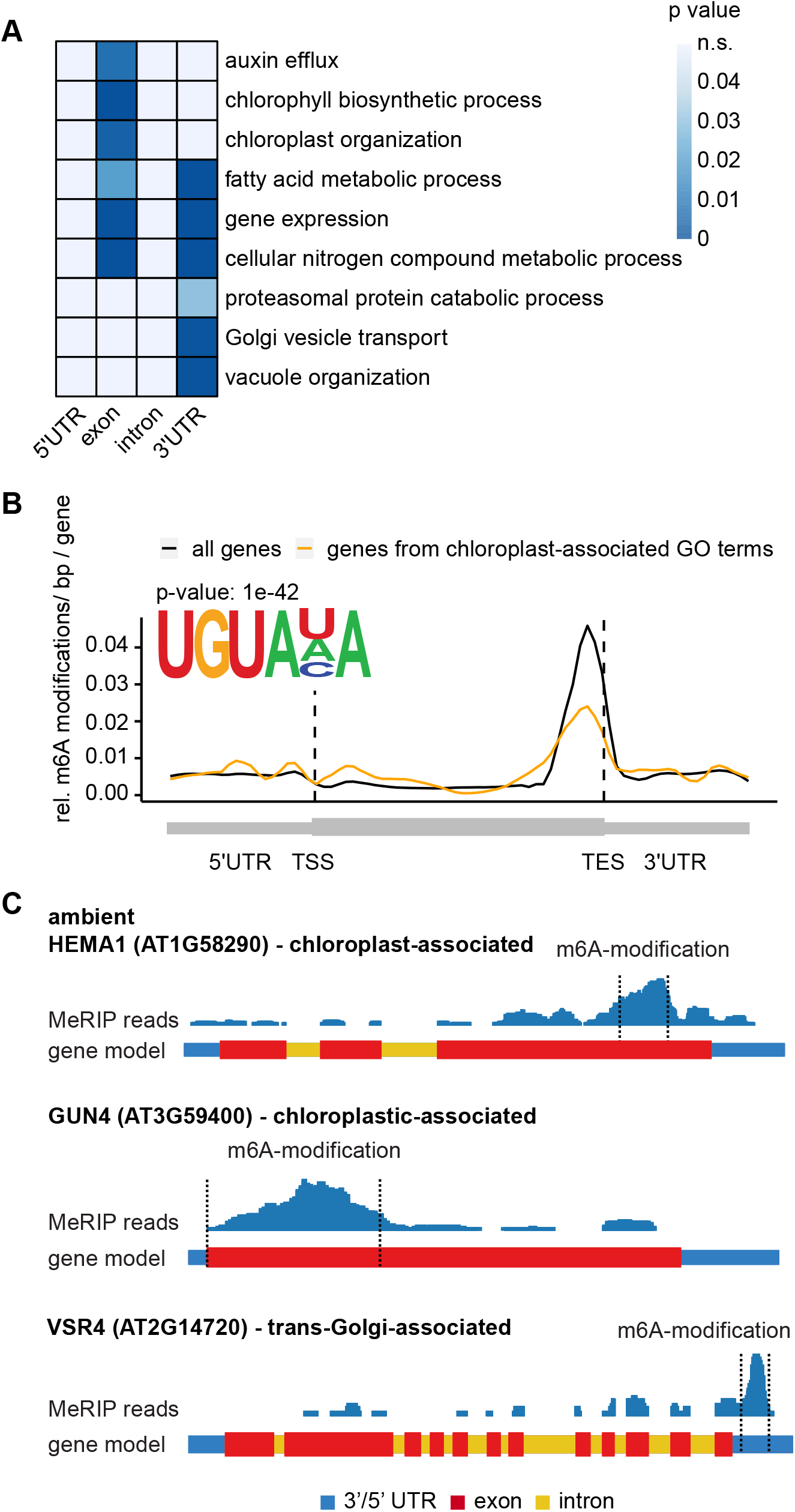
Region of m^6^A modification on transcripts might predict m^6^A function. **(A)** Heatmap of Gene Ontology (GO) term enrichment for transcripts harboring m6A-modifications in 3’UTR, exon, intron, and 5’UTR regions. Heatmap colors represent p-values of associated GO terms. Data from all samples was combined. Chloroplast-associated GO terms were enriched for genes that harbor m6A transcript modifications in exon regions and endomembrane system-associated GO terms were enriched for genes that harbor m6A transcript modifications in 3’UTRs. **(B)** Metagene analysis showing MeRIP-seq read enrichment along transcripts. Sequences 5000 bp upstream of the transcription start site (TSS) and 5000 bp downstream of the transcription end site (TES) were considered in the analysis. Data from all samples was combined. Genes associated with chloroplast GO terms (orange) are compared to all genes (black). m6A-containing sites for chloroplast GO terms were analyzed for enriched RNA sequences. The highest-ranking motif is shown together with the corresponding p-value. Chloroplast-associated genes show less m6A modification close to the 3’UTR when compared to all other genes. **(C)** Gene plots for *HEMA1, GUN4*, and *VSR4*. Normalized coverage of MeRIP-sequencing reads from plants grown in ambient conditions is shown in blue. Identified m^6^A-modification of the transcripts is indicated by dashed lines.

We further analyzed the location of m^6^A-modifications in exons of genes that were associated with chloroplasts. To this end, we performed metagene and motif analyses. Comparing the distribution of m^6^A-modifications along the gene body between all genes and chloroplast-associated genes, we identified fewer transcripts that are modified close to the TES (approx. 0.02 m^6^A-modifications / bp / gene vs. approx. 0.05 m^6^A-modifications / bp / gene) (Figure 4B, C). We also observed a higher relative number of m^6^A-modifications in the beginning of coding regions of genes. Examples of transcripts involved in chloroplast or trans-Golgi-associated processes and their m^6^A-modification are shown in Figure 4C. The combined results suggest that m^6^A-modification of transcripts might be important for transcript or protein localization. According to our observations m^6^A-modification of transcript close to the TES might inhibit chloroplast localization. Similarly, transcript m^6^A-modification in 3’UTRs might be important for localization to the endomembrane system such as the trans-Golgi network or the vacuole.

### The environment controls differential m^6^A-modification of transcripts

We next investigated if environmental conditions led to differential m^6^A-modification of transcripts by performing GO term analyses on exclusively m^6^A methylated transcripts for each condition (Fig 5A). The GO terms seed development and reproduction were significantly enriched in shade and HT. Therefore, m^6^A-modification of those transcripts might be a common regulatory mechanism to adapt to shade or HT conditions. In contrast, GO terms generation of precursor metabolites and catabolic process were significantly enriched in ambient conditions but not in shade nor in HT, which indicates that the respective treatments might lead to specific methylation in ambient or demethylation in treated conditions. In HT, we also find transcripts of genes associated with thylakoid membrane organization and RNA processing to be enriched. When compared to HT, ambient plants exhibited an enrichment in transcripts associated with responding to cold. The molecular mechanisms underlying those adaptions are not fully understood and it is feasible that m^6^A could play a key role. When comparing etiolated and de-etiolated seedlings, we observed transcripts associated with cell growth, plant organ morphogenesis, and cell differentiation to be significantly enriched. Etiolated seedlings exhibit elongated hypocotyls, which are mediated by cell elongation and differentiation. Upon de-etiolation, plants fully develop their photosynthetic apparatus. Interestingly, GO terms related to chlorophyll biosynthesis, photosynthesis or photomorphogenesis were exclusively enriched in de-etiolated plants suggesting that m^6^A-modification plays an important role in those adaptive responses.

**Figure 5.**
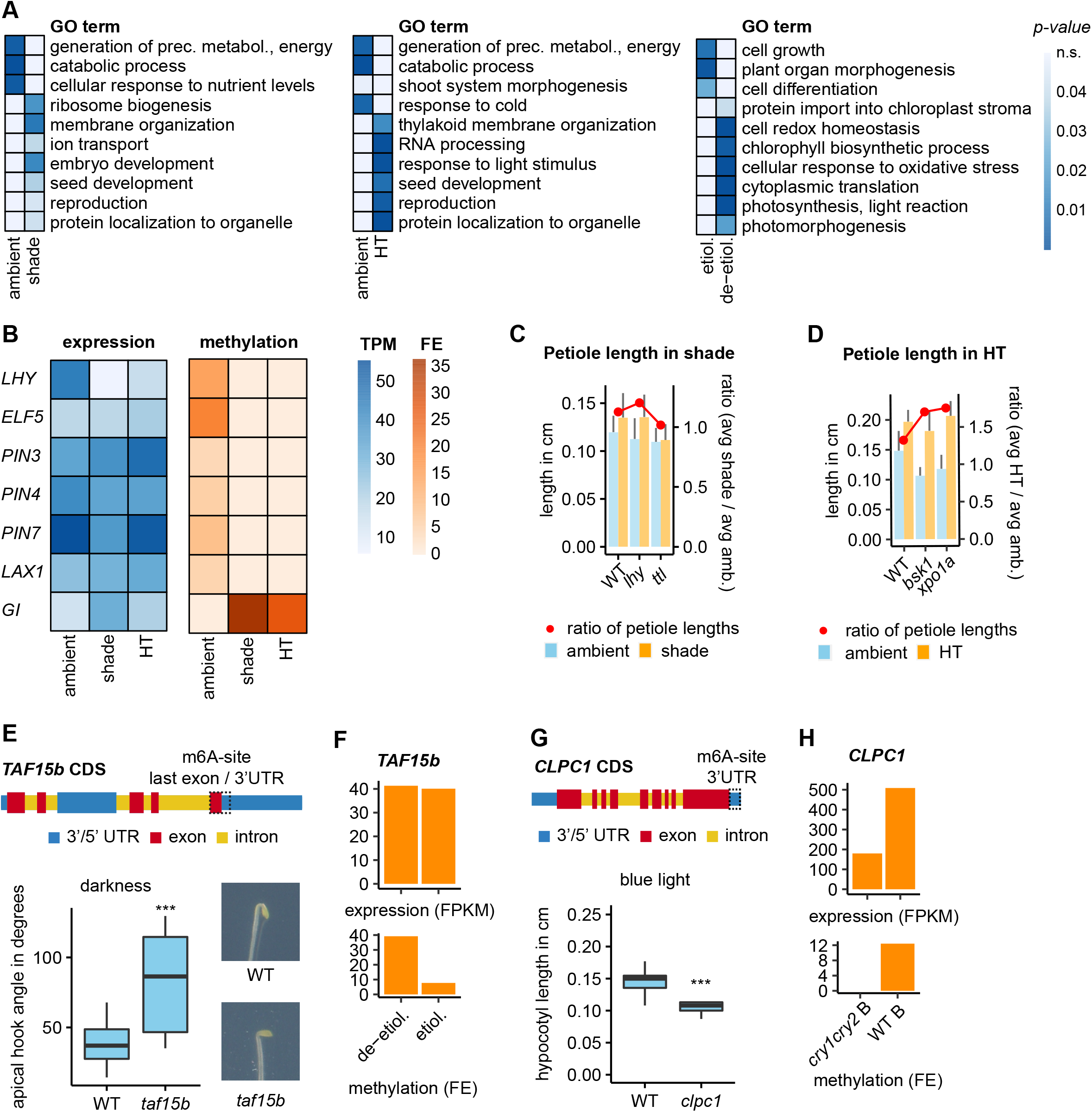
Differential methylation might be necessary for adaptive responses. **(A)** Heatmap of GO term enrichment for transcripts harboring m^6^A-modifications in ambient vs. shade (left), ambient vs. high temperature (HT) (middle), and etiolated vs. de-etiolated (right). Heatmap colors represent p-values of associated GO terms. Distinct groups of GO terms are enriched for specific conditions. **(B)** Heatmaps of transcript abundance (TPM in blue) and methylation (fold enrichment, FE in orange) between plants grown in ambient (left), shade (middle) or high-temperature (HT, right) conditions. Gene expression does not correlate with methylation. **(C)** Petiole length of plants grown in shade conditions are shown as bar plot. Error bars represent standard deviation. Red dots represent the elongation response as calculated by the ratio of the average length in shade divided by the average length in ambient condition (N ≥ 14). When grown in shade, *lhy* mutant lines exhibited longer petioles compared to WT, while *ttl* mutant lines exhibited shorter petioles than WT. **(D)** Petiole length of plants grown in HT conditions are shown as bar plot. Error bars represent standard deviation. Red dots represent the elongation response as calculated by the ratio of the average length in HT divided by the average length in ambient condition (N ≥ 14). *bsk1* and *xpo1a* mutant lines exhibited longer petioles than WT when grown in HT. **(E)** The gene model for *TAF15b* is represented by colored boxes. The identified site of m^6^A modification is marked with a dashed box. Apical hook angles of dark-grown seedlings were measured. Asterisks indicate significant differences as determined by Student’s t-test (***: *p* < 0.0005) between WT and *taf15b* plants (N = 13). Pictures of representative seedlings are shown. *taf15b* mutant lines have partially open apical hooks when grown in darkness. **(F)** Expression (FPKM) and methylation (fold enrichment, FE) are shown for *TAF15b* in etiolated/de-etiolated conditions. *TAF15* transcripts show stronger methylation in de-etiolated seedlings but no significantly different abundance. **(G)** The gene model for *CLPC1* is represented by colored boxes. The identified site of m^6^A modification is marked with a dashed box. Hypocotyl length in blue light was measured. Asterisks indicate significant differences as determined by Student’s t-test (***: *p* < 0.0005) between WT and *clcp1* plants (N ≥ 13). *clpc1* mutant lines exhibit significantly shorter hypocotyl lengths in blue light compared to WT. **(H)** Expression (FPKM) and methylation (fold enrichment, FE) are shown for *CLPC1* in WT and *cry1cry2* plants grown in blue light. *CLPC1* transcripts are differentially methylated and differentially expressed.

We further analyzed specific genes with differentially m^6^A-modified transcripts including genes involved in developmental adaptive responses through the circadian clock or auxin signaling, which show highly specific m^6^A-modification patterns (Figure 5B). *LHY* (*LATE ELONGATED HYPOCOTYL*), *ELF5* (*EARLY FLOWERING 5*), *PIN3, PIN4, PIN7* (*PIN-FORMED3, 4, 7*), and *LAX1* (*LIKE AUXIN RESISTANT 1*) were m^6^A-modified exclusively in ambient conditions and not in shade or HT conditions. The opposite is true for *GI* (*GIGANTEA*), which was exclusively m^6^A-modified in shade and HT, but not in ambient conditions. Interestingly, there was no correlation between differential expression and differential m^6^A methylation which underlines previous observations suggesting that m^6^A does not function exclusively by regulating mRNA abundance. Components of the circadian clock, in particular *LHY*, plays a key role in adaptive processes to shade (Wang et al., 2016) and for temperature compensation of the circadian clock (Gould et al., 2006). PIN proteins are essential for hypocotyl elongation and shade avoidance response. Plants lacking functional PIN3 do not exhibit a shade avoidance response (Keuskamp et al., 2010) and *pin3, pin4, pin7* triple mutant seedlings exhibit significantly reduced hypocotyl elongation in response to elevated temperatures (Bellstaedt et al., 2019). The observed differential methylation of those genes might therefore be functionally relevant to adapt to environmental conditions.

To find new genes that might be involved in adaptive responses, we exposed mutant lines of genes with differentially m^6^A-modified transcripts to different environmental conditions. In shade, we observed that *lhy* mutant plants showed an approximately 8% stronger induction of petiole elongation compared to WT (Figure 5C). In our dataset transcripts of *LHY* were exclusively m^6^A-modified in ambient, but not in shade conditions (Figure S5A). In contrast to that, *Arabidopsis* lines lacking functional TTL (TRANSTHYRETIN-LIKE PROTEIN) lacked the petiole elongation response when grown in shade. *TTL* is part of the brassinosteroid pathway, but has not been connected to the shade avoidance response. *TTL* mRNA is m^6^A-modified in ambient, but not shade conditions (Figure S5A). Neither *lhy* nor *ttl* mutant plants exhibited significant alterations in their hypocotyl elongation response indicating that both genes exclusively regulate petiole elongation (Figure S5B). In HT, we found *bsk1 (brassinosteroid-signaling kinase 1)* and *xpo1a (Arabidopsis thaliana exportin 1)* mutant lines to exhibit enhanced petiole elongation (1.7- and 1.8-fold elongation in HT compared to ambient) when compared to WT (1.3-fold elongation in HT compared to ambient) (Figure 5D). *BSK1* transcripts are m^6^A-modified in ambient, but not HT conditions, while *XPO1A* transcripts are modified exclusively in HT conditions (Figure S5A). Regarding HT-induced hypocotyl elongation, *bsk1* but not *xpo1a* mutant lines showed an enhanced response (1.8 and 1.3-fold elongation in HT compared to ambient) when compared to WT (1.3-fold elongation in HT compared to ambient) (Figure S5C). Interestingly, the brassinosteroid pathway is important for thermomorphogenic adaptions, but previous studies have mainly focused on the role of *BZR1* (*BRASSINAZOLE RESISTANT 1*) (Ibañez et al., 2018). XPO1A is essential for plant heat tolerance, but its function in adaptions to elevated non-stress heat conditions has not been described (Wu et al., 2010).

We further investigated other adaptive responses such as apical hook formation in etiolated seedlings or hypocotyl growth inhibition in blue light (Figure 5E-H). *TAF15b* (*TATA-BINDING PROTEIN-ASSOCIATED FACTOR 15b*) is involved in flowering time control (Eom et al., 2018). The transcript of *TAF15b* is m^6^A-modified in close proximity to the TES almost exclusively in de-etiolated seedlings (5-fold increase in FE), while transcript levels are not significantly different between etiolated or de-etiolated seedlings (Figure 5E, F). Seedlings without functional *TAF15b* exhibit partial apical hook opening when grown in darkness. Hence, *TAF15b* is essential for apical hook development in darkness and *TAF15b* is not regulated through differential gene expression. It is feasible that *TAF15b* function is regulated through differential m^6^A-modification. Similarly, we observed transcripts of *CLPC1* (*CASEINOLYTIC PROTEASE COMPLEX 1*) to be m^6^A-modified in the 3’UTR region (Figure 5G). The transcript levels are significantly different between WT and *cry1cry2* mutant plants grown in blue light, but high in both samples (FPKM > 170), yet we identified m^6^A-modified transcripts exclusively in WT plants. Our analysis of hypocotyl lengths in blue light showed a significantly reduced hypocotyl length in *clpc1* mutants when compared to WT plants. It is possible that *CLPC1* m^6^A-modification leads to a stabilization of *CLPC1* transcripts in blue light, which is important to prevent a hypersensitive light response.

### m^6^A might affect gene function by controlling alternative polyadenylation

In humans, the UGUA motif serves as a binding domain for the Human Cleavage Factor I (CFI_m_) complex (Yang et al., 2011). The CFI_m_ complex is part of the pre-mRNA 3’ end processing apparatus and important to regulate 3’UTR length by promoting usage of distal APA sites, thus producing longer 3’UTRs (Martin et al., 2012). APA is implicated in mRNA stability, mRNA localization, and miRNA binding (Mayr, 2017). Interestingly, m^6^A-modification of 3’UTR regions has also been associated with APA with methylated transcripts showing a higher preference for proximal polyadenylation sites (Luo et al., 2020; Parker et al., 2020; Yue et al., 2018). To test whether m^6^A might regulate APA in the context of adapting to varying environmental conditions, we analyzed our RNA-seq dataset (Xia et al., 2014) to identify APA events associated with m^6^A deposition using the DaPars algorithm (Xia et al., 2014). Depending on the respective condition, approximately 26 – 81% of methylated transcripts exhibit APA events (Figure 6A). *cry1cry2* mutant plants grown in blue light showed the least amount of overlap (26%) between APA and m^6^A, whereas etiolated plants showed the biggest overlap (81%). Hence, we identified a large set of genes that are m^6^A-modified and show APA suggesting a functional connection.

**Figure 6.**
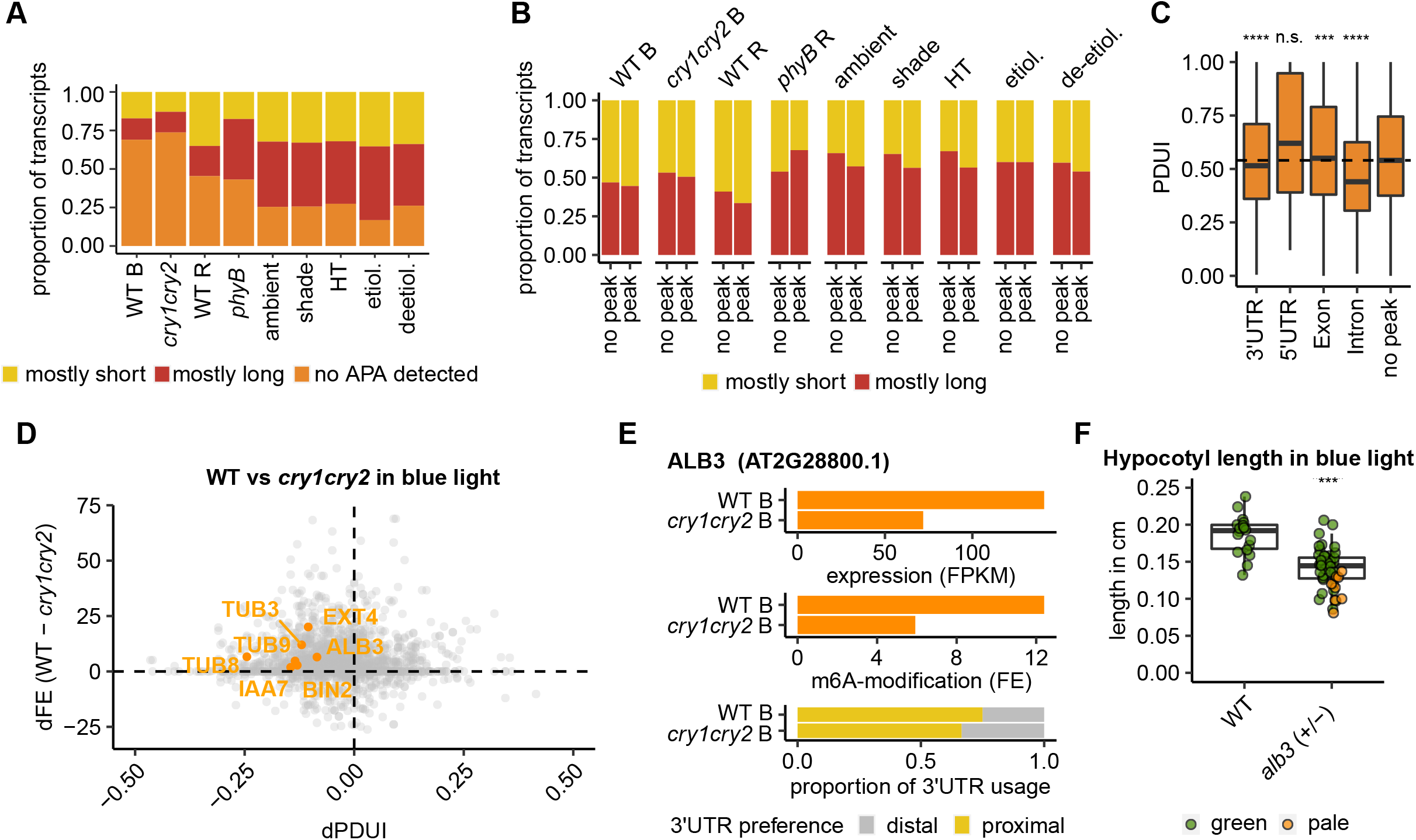
m^6^A influences transcript abundance via APA. **(A)** Preferred 3’UTR length of transcripts that were identified to be m^6^A-modified and show APA. PDUI < 0.5: mostly short 3’UTR, PDUI > 0.5: mostly long 3’UTR. A large number of m^6^A-modified transcripts exhibit APA. **(B)** Comparison of APA between genes with or without detected m^6^A-modification. PDUI < 0.5: mostly short 3’UTR, PDUI > 0.5: mostly long 3’UTR. m^6^A-modified transcripts are shorter than unmodified transcripts in most conditions. In *phyB* grown in red light, m^6^A-modified transcripts are longer and in etiolated seedlings no difference between m^6^A-modified and unmodified transcripts was observed. **(C)** Combined PDUI of transcripts harboring m^6^A-modifications for specific regions of the transcript from all samples. The dotted line depicts median PDUI for genes that were not identified to be m^6^A-modified (no peak). Asterisks indicate significant differences as determined by Student’s t-test (***: p < 0.001, ****: *p* < 0.0001) between samples with no peaks and the respective region. Transcripts with m^6^A modifications in 3’UTRs or introns are shorter than unmodified transcripts, while transcripts with m^6^A-modifications in exons are shorter. **(D)** The difference in 3’UTR usage preference (dPDUI) is compared to the difference in m^6^A fold enrichment (dFE) for each gene in WT and *cry1cry2* plants treated with blue light. The transcripts of a large number of genes are shorter and more methylated in WT. **(E)** Direct comparison of *ALB3* expression (FPKM), methylation (fold enrichment, FE), and 3’UTR usage preference. Transcript abundance, m^6^A modification and 3’UTR usage are correlated for *ALB3*. **(F)** Hypocotyl length in blue light was measured. Asterisks indicate significant differences as determined by Student’s t-test (***: *p* < 0.0005) between WT and pale *alb3* plants (N = 8). Pale *alb3* mutant seedlings exhibit significantly reduced hypocotyl lengths compared to WT when grown in blue light.

To further investigate the effect of m^6^A on APA, we compared the preference for proximal or distal polyadenylation sites in transcripts with or without detected m^6^A-modifications. The preference for either polyadenylation site is measured by the “percentage of distal polyadenylation site usage index” (PDUI). A PDUI value above 0.5 indicates a preference for distal polyadenylation sites. Conversely, a PDUI value below 0.5 indicates a preference for proximal polyadenylation sites. m^6^A-modified transcripts showed a higher preference for proximal polyadenylation sites (PDUI < 0.5) than non-m^6^A-modified transcripts, thus producing shorter 3’UTRs (Figure 6B, S6A). Notably, m^6^A-modified transcripts from *phyB* mutant seedlings grown in red light had longer 3’UTRs when compared to non-m^6^A-modified transcripts. Etiolated plants did not exhibit a difference in 3’UTR length preference between transcripts with or without m^6^A-modification. These results suggest that under most conditions, m^6^A-modification of RNA leads to a preference for short 3’UTRs. In red light, this process might be regulated by PHYB, since the response is reversed in *phyB* mutant plants. After combining all m^6^A-sites and the PDUI of respective transcripts our analysis suggests, that the preference for shorter 3’UTRs is mainly driven by the deposition of m^6^A in 3’UTR (median PDUI: 0.515) or intron (median PDUI: 0.44) regions of transcripts when compared to non-m^6^A-modified transcripts (median PDUI: 0.54) (Figure 6C). Interestingly, transcripts with m^6^A-modifications in their 5’UTR seem to mainly use distal polyadenylation sites (median PDUI: 0.62). This suggests that m^6^A-modified 3’UTR regions promote usage of proximal polyadenylation sites and thus shorter 3’UTRs, while m^6^A-modified 5’UTR regions might promote the usage of distal polyadenylation sites.

A more detailed analysis of plants grown in blue light revealed that 49% of transcripts with a predicted APA event show stronger m^6^A-modification as defined by a higher fold enrichment of reads between input and MeRIP in WT compared to *cry1cry2* mutant lines grown in blue light (dFE). Moreover, a subgroup of those m^6^A-modified transcripts exhibits a strongly increased preference for proximal polyadenylation sites in WT as measured by the difference in PDUI between the two samples (dPDUI) (Figure 6D). In this group, we find a host of genes that are necessary for hypocotyl elongation such as *EXTENSIN 4* (*EXT4*) or *TUBULIN 3, 8*, and *9* (*TUB3, 8, 9*) as well as genes involved in hormone signaling such as *INDOLE-3-ACETIC ACID 17* (*IAA17*), and *BRASSINOSTEROID-INSENSITIVE 2* (*BIN2*). Additionally, a gene important for proper chloroplast development, *ALBINO 3* (*ALB3*), is suggested to be regulated via m^6^A-controlled APA (Figure 6E, Figure S6B). ALB3 is essential for chloroplast development as evidenced by *alb3* mutants that develop white pale cotyledons and do not survive on soil past the seedling stage (Sundberg et al., 1997). Hence, *ALB3* is important for adapting to surrounding light conditions and optimization of chloroplast development. *ALB3* transcript abundance is 2-fold upregulated in WT when compared to *cry1cry2* mutant plants. While transcripts are m^6^A-modified in both genotypes, the fold enrichment in WT plants is approximately two times higher than in *cry1cry2* mutants. Additionally, the usage of a proximal 3’UTR polyadenylation site is 9% higher in WT plants. It is conceivable that the CRYs mediate proper m^6^A-modification of *ALB3* transcripts. In the absence of *CRY*s, *ALB3* transcripts are less methylated, which in turn might lead to a preference for shorter 3’UTRs, thus stabilizing the transcripts. This might explain part of the higher *ALB3* transcript abundance in WT when compared to *cry1cry2* mutants grown in blue light. To further investigate a possible role of ALB3 in blue light signaling, we analyzed *alb3* mutants grown in blue light (Figure 6F), and found that their hypocotyls were significantly shorter in blue light compared to WT. This data suggests that *ALB3* might play an important role in mediating hypocotyl growth in blue light conditions. In summary, we conclude that CRY-mediated blue light signaling might influence gene expression by regulating APA through differential m^6^A-modification of transcripts to finetune adaptive responses.

Taken together our data suggest that environmental stimuli, particularly blue light, control RNA m^6^A-modifications which in turn influence APA to finetune basic biological functions such as hypocotyl elongation, greening, and blue light signaling. This effect does not seem to be universal but specific to certain transcripts. Moreover, we find that methylation of specific regions of transcripts are associated with their localization or localization of encoded proteins. Further experiments will be necessary to pinpoint the molecular mechanisms underlying these processes and if APA and mRNA localization are connected through differential m^6^A-modification.

## DISCUSSION

Despite advances in our understanding of m^6^A-modifications and their effect on gene regulatory mechanisms, our knowledge of how environmental signals shape the m^6^A landscape and elicit transcriptional changes remains limited (Anderson et al., 2018; Arribas-Hernandez et al., 2018; Li et al., 2014; Scutenaire et al., 2018). Here, we focused on how the light and temperature environment influences m^6^A-modifications to better understand the effect of individual stimuli on the m^6^A RNA landscape in *Arabidopsis*. Light and temperature are crucial environmental signals for plants and control various processes throughout plants’ life cycles (Casal and Balasubramanian, 2019; Neff and Chory, 1998). We provide first evidence for the role of light and temperature in controlling m^6^A RNA modifications and propose that influencing transcript/protein localization and APA are possible downstream mechanisms for gene regulation in this context. We used MeRIP-seq to examine the transcriptome-wide m^6^A landscape and found that plants exhibit a highly specific m^6^A RNA signature depending on their environment, enabling discovery of new genes in these pathways. We were able to identify a large number of transcripts that were m^6^A-modified under varying light and temperature conditions, and the number of identified m^6^A-modified transcripts varied strongly between conditions. MeRIP assays were conducted under the same conditions and replicated, suggesting the differences in m^6^A RNA abundance are physiologically relevant and not indicative of technical issues. Previous reports have provided evidence that in mice global m^6^A RNA abundance in specific tissue types can be regulated by the circadian clock or stress (Engel et al., 2018; Wang et al., 2015). Similarly, we found that the environment might induce global changes in m^6^A RNA abundance in plants. Notably, plants grown in red light and darkness (etiolated) exhibited the strongest reduction in identified m^6^A-modified transcripts. It is conceivable that red light downregulates the activity of methylases or upregulates the activity of demethylases. However, in our RNAseq datasets we were unable to identify any significant upregulation nor downregulation of known m^6^A eraser and writer genes respectively in seedlings grown in red light. Taking into consideration that WT and *phyB* mutant plants showed a reduced number of m^6^A-modified transcripts, it is likely that this process is not mediated by *phyB*. There may be uncharacterised m6A methylases or demethylases which mediate m^6^A under red light. Alternatively, components of the m^6^A machinery may be regulated post-translationally in red light.

We found regions around the TES and 5’UTR to be highly m^6^A-modified, in line with previous observations in *Arabidopsis* (Luo et al., 2014). While 5’UTR m^6^A-modification is not highly enriched in metazoans, m^6^A-sites in the 5’UTR have been identified and functionally associated with promoting cap-independent translation (Meyer et al., 2015). It will be worth investigating whether m^6^A-modification of 5’UTR regions has a similar effect in *Arabidopsis* and to understand why plants have evolved a higher rate of m^6^A-modification at the 5’UTR than metazoans. The observation that certain regions of transcripts might be more or less methylated under different conditions could be a way of fine tuning the transcriptome on a more global scale. The m^6^A methyltransferase component VIR-LIKE M^6^A METHYLTRANSFERASE ASSOCIATED (VIRMA) mediates the preferential m^6^A-modification of 3’UTRs in HeLa cells (Yue et al., 2018) and thus provides evidence that the regulation of VIRMA might lead to global effects on 3’UTR m^6^A-modification. So far, a region-specific methylation factor in plants has not been discovered.

Motif analysis revealed a strongly enriched UGUA motif in 3’UTR regions that were identified to be m^6^A-modified. Currently, there is an ongoing debate about the importance of UGUA motifs in *Arabidopsis* m^6^A-modified transcripts: Studies have reported the presence of the UGUA motif in m^6^A-modified regions (Miao et al., 2019) and URUA has been described as the main binding motif for the m^6^A reader protein EVOLUTIONARILY CONSERVED C-TERMINAL REGION 2 (ECT2) (Wei et al., 2018). However, other studies find the RRACH consensus motif, identified in metazoans, to be highly enriched in *Arabidopsis* m^6^A-modified transcripts (Anderson et al., 2018; Luo et al., 2014; Parker et al., 2020; Shen et al., 2016). It has been suggested that UGUA motifs are in spatial proximity to m^6^A-modified regions, but might not themselves be m^6^A-modified. In the deep learning approach, we took sequences 200 bp up- and downstream of the identified m^6^A-site into account, but did not identify an enrichment of the RRACH motif. This suggests that UGUA might indeed be directly modified. Single-nucleotide mapping methods for m^6^A-modification of *Arabidopsis* RNA will be necessary to conclusively define motifs for m^6^A-modifications. After binning identified m^6^A-modification into different regions of transcripts, such as 5’UTR, exon, intron, and 3’UTR, we found the UGUA sequence to be enriched in peaks within 3’UTRs, but not other regions. Interestingly, for all other regions, motifs were identified that contained a predominant adenosine (A), which is likely to be methylated. We hypothesize that transcripts are m^6^A-modified within various sequence contexts, which controls the modification of specific regions of transcripts. Previous analyses might have missed this observation because they focused on global m^6^A-modification sites. In this scenario, the dominance of peaks in 3’UTRs might mask the presence of alternative motifs in other regions. Note that p-values for non-3’UTR regions are lower than for the 3’UTR. This might be due to the lower number of peaks in those regions. Similarly, the low abundance of identified m^6^A-modifications in WT and *phyB* mutant plants grown in red light, are likely the reason for the comparatively low p-value associated with the identified RNA motif sequence.

Our observation that the m^6^A-modification of exons correlates with transcripts of genes that are associated with chloroplast-related GO terms could be relevant for the localization of those transcripts. Certain transcripts encoding proteins that are specific to the ER, chloroplasts or mitochondria preferentially localize to organellar membranes (Tian et al., 2019). To date, the underlying molecular mechanisms for targeted mRNA transport are not well understood. It has been suggested that cis-elements in the 3’UTR of *VDAC3* control its mRNA localization to mitochondria (Michaud et al., 2014b). Since m^6^A has been reported to influence APA and therefore 3’UTR lengths in plants (Parker et al., 2020), tightly regulated m^6^A-modification might lead to the inclusion or exclusion of RNA zip codes that are important for mRNA localization. Moreover, m^6^A-binding proteins, called YTH-domain proteins, are important for mRNA localization in mammalian cells (Madugalle et al., 2020; Wang et al., 2014). Specific m^6^A-modification of transcripts could influence the binding of YTH-domain proteins to mRNA and thus mRNA localization. m^6^A-modifications at specific sites could therefore impair or promote proper mRNA localization controlling efficient protein translation and organellar import, thus influencing development.

GO term analyses of genes with methylated transcripts between tested conditions revealed that transcripts of genes involved in various housekeeping processes are not differentially methylated. However, we found GO terms of certain biological processes to be significantly enriched only under specific conditions. For example, exclusively in HT, transcripts of genes that are involved in RNA processing are highly enriched. RNA processing, particularly mRNA alternative splicing has long been recognized to be regulated by temperature (Casal and Balasubramanian, 2019; Lin and Zhu, 2021). Moreover, m^6^A plays a role in alternative splicing (Adhikari et al., 2016; Meyer and Jaffrey, 2017). The immediate molecular connection between m^6^A-modifications and alternative splicing in high temperature has not been elucidated yet. It is noteworthy that chloroplasts are very sensitive to heat and plants have evolved intricate mechanisms to protect chloroplasts from the negative effects of high temperatures (Hu et al., 2020). Since in our dataset a large set of nuclear-encoded chloroplast genes were identified to be m^6^A-modfied it is feasible that m^6^A-modification may act as an important regulatory mechanism to fine-tune temperature responses.

We identified new candidate genes in pathways of adaptive developmental responses, such as blue light-mediated photomorphogenesis, skotomorphogenesis, shade avoidance, and thermomorphogenesis by investigating differential m^6^A-modification of transcripts. For example, *TAF15b* has been described to play a role in the autonomous flowering pathway (Eom et al., 2018) and autoimmunity (Dong et al., 2016). Our data suggest a so far undescribed role of *TAF15b* in promoting apical hook formation in etiolated seedlings. Additionally, we found that CLPC1 promotes hypocotyl elongation in blue light. In the past, *CLPC1* has been associated with disease resistance and thermotolerance (Pant et al., 2020), but not blue light signaling. Since CLPC1 is mainly localized in the chloroplast, it is interesting to speculate about retrograde signaling mechanisms. Recent reports have demonstrated that light and plastid signaling are strongly connected (HernándezLVerdeja et al., 2020) and it will be worth investigating whether *CLPC1* contributes to this signaling pathway. Similarly, we have found other genes involved in shade and HT responses, that should be investigated further.

In metazoans, the UGUA motif is an important binding motif for the Human Cleavage Factor I (CFI_m_) complex (Yang et al., 2011). Upon binding to RNA, this complex mediates the distal usage of APA sites, thus promoting longer 3’UTR regions (Martin et al., 2012). Moreover, there is a direct connection of m^6^A-modification in the 3’UTR and APA: Transcripts with an m^6^A-modification exhibit a strong bias towards using more proximal APA sites (Molinie et al., 2016) and VIRMA directly associates with polyadenylation cleavage factors to alter APA (Yue et al., 2018). Depletion of VIRMA leads to an increased use of distal polyadenylation sites. In contrast to that, it was reported that m^6^A inhibits the use of proximal APA sites in plants (Parker et al., 2020). The m^6^A methyltransferase component FKBP-12 INTERACTING PROTEIN 37 (FIP37) and the m^6^A-binding protein CLEAVAGE AND POLYADENYLATION SPECIFICITY FACTOR COMPLEX 30L (CPSF30L) mediate alternative polyadenylation (Pontier et al., 2019). These findings prompted us to further investigate whether the environment influences m^6^A-modifications, thus influencing APA as a possible mechanism for plants to adapt to their surrounding light and temperature conditions. Indeed, in our dataset we identified a large number of transcripts that show APA events and are m^6^A-modified. Since m^6^A-modified transcripts exhibited a bias towards a higher usage of proximal polyadenylation sites, we contribute to the ongoing discussion about the effect of m^6^A on APA by providing evidence that m^6^A might directly shift the usage of polyadenylation sites towards more proximal sites and does not inhibit proximal APA site usage.

To elucidate the physiological relevance of the shifting 3’UTR length preference, we focused further analyses on WT and *cry1cry2* mutant seedlings grown in blue light. We found a number of genes that are responsible for adaptation to and growth in blue light being differentially m^6^A-modified and showing a shift towards shorter 3’UTRs between the two genotypes. In the case of ALB3 this observation is accompanied with a higher transcript expression. It is feasible that transcripts are regulated through m^6^A-mediated APA. To date, various stabilizing and destabilizing effects of m^6^A on transcripts have been described (Anderson et al., 2018; Martinez-Perez et al., 2017; Parker et al., 2020; Shen et al., 2016; Wei et al., 2018). Here, we provide evidence for how the environment might influence m^6^A-mediated transcript stabilization via APA. Since m^6^A-modifications seem to have highly context-dependent and opposing outcomes, such as stabilization or destabilization of transcripts, the m^6^A-mediated transcript regulation via APA is most likely one of many effects m^6^A has on gene regulation. We present one possible way for a downstream regulatory mechanism, but more studies will be necessary to fully elucidate the complex role of m^6^A in this regard. As discussed above, it would also be feasible that APA leads to the inclusion or exclusion of cis-elements, or zip codes, regulating localization of transcripts. Since ALB3 is mainly localized in the chloroplast, potentially impairing mRNA localization might have a strong effect on protein abundance in the chloroplast.

In conclusion, our data suggest that the environment significantly shapes the m^6^A RNA landscape in plants. Differential m^6^A-modification of transcripts might influence APA to fine tune the abundance and localization of specific transcripts. In the future it will be interesting to elucidate the molecular mechanisms governing these adaptive responses.

## Acknowledgements

This work is supported by National Institutes of Health (NIH) grant R35GM125003 to U.V.P. Figure 1A was created using BioRender.com. We thank D. Jackson, U. Hoecker, Y. Hu for critical reading of this manuscript.

## Materials and Methods

### Plant materials and growth conditions

The *Arabidopsis thaliana* accession Col-0 (WT) was used as WT for all experiments. *cry1-304 cry2-1* (Mockler et al., 1999) and *phyB-9* (Reed et al., 1993) were used as photoreceptor mutants. The following T-DNA insertion lines were obtained from the Arabidopsis Biological Resource Center (ABRC): *alb3* (AT2G28800, CS16), *bsk1* (AT4G35230, SALK_122120), *clpc1* (AT5G50920, SALK_014058C), *lhy* (AT1G01060, CS9379), *taf15b* (AT5G58470, SALK_061974C), *ttl* (AT5G58220, SALK_137289), *xpo1a* (AT5G17020, SALK_028886) and genotyped by PCR to obtain homozygous lines. Seeds were sown on 0.5x Linsmaier Skoog basal medium (Himedia Laboratories) solidified using 0.8% agar and stratified for 2 days at 4°C. Germination was initiated by exposing them to white light for 1.5 – 3h. Seedlings for total RNA isolation were treated as follows: WT and *cry1-304 cry2-1* plants were grown in continuous blue light (10 μmol m^-2^ s^-1^) for 4 days. WT and *phyB-9* plants were grown in continuous red light (20 μmol m^-2^ s^-1^) for 4 days. Monochromatic light conditions were established by an LED light source (Percival). For the de-etiolation assay, WT plants were grown in darkness for 4 days and either incubated in white light (95 μmol m^-2^ s^-1^) or kept in darkness for additional 24 h. For the second set, WT plants were grown in continuous white light (95 μmol m^-2^ s^-1^, red:far-red ratio = 5.4, 22°C) for 4 days. Subsequently, plants were kept in these conditions for 24 h or subjected to either shade (70 μmol m^-2^ s^-1^, red:far-red ratio = 0.2, 22°C) or high temperature (95 μmol m^-2^ s^-1^, red:far-red ratio = 5.4, 30°C) conditions for the same 24 h time period. To obtain enough plant material for total RNA isolation, tissue from 10-15 plates was pooled for each condition and replicate. Tissue was snap-frozen in liquid nitrogen until further processing.

### Phenotypic analysis

Plants were sown, stratified, and germinated as above. Phenotyping was performed under the following conditions. For assessment of apical hooks, plants were wrapped in three layers of aluminum foil and grown in darkness for 4 d. Phenotyping in blue light was performed by exposing seedlings to 10 µmol m^-2^ s^-1^ constant blue light for 4 d. For shade treatment, plants were first grown in constant white light (100 μmol m^-2^ s^-1^, red/far-red ratio = 8, 22°C) for 3 d and either kept in those conditions for additional 4 d or subjected to shade conditions (60 µmol m^-2^ s^-1^, red/far-red ratio = 0.4, 22°C) for 4 d. Similarly, plants for high temperature treatments were grown in constant white light (100 μmol m^-2^ s^-1^, 22°C) for 3 d and either kept in those conditions for additional 4 d or subjected to high temperature conditions (100 µmol m^-2^ s^-1^, 30°C) for 4 d. After the treatment, plates were scanned using a flatbed scanner and metrics such as apical hook angle, hypocotyl length or petiole length were determined using Fiji (Schindelin et al., 2012).

### RNA isolation

Total RNA was isolated following a protocol modified from (Chomczynski and Sacchi, 2006). In brief, plant tissue was ground to a fine powder using pestle and mortar. Approximately 0.5 g of ground tissue was incubated with 5 ml of acid guanidinium thiocyanate-phenol-chloroform reagent (0.8 M guanidinium thiocyanate, 0.4 M ammonium thiocyanate, 0.1 M sodium acetate pH 5.0-5.2, 5% glycerol, 38% (v/w) phenol) at room temperature for 5 min. 1.5 ml chloroform was added, mixed by inversion and incubated at room temperature for 3 min. Samples were centrifuged at 12,000 x g and 4°C for 10 min. After phase separation, the aqueous phase was transferred to a new tube and 2.5 ml of isopropanol was added, mixed and incubated at room temperature for 10 min. Samples were centrifuged at 12,000 x g and 4°C for 10 min. The supernatant was discarded and the pellet was washed in 1 ml of 75% ethanol and incubated at -20°C for 10 min. Afterwards samples were centrifuged at 12,000 x g and 4°C for 5 min and the supernatant removed. This washing step was performed again for a total of 2 washes. The RNA pellet was resuspended in 30-50 μl of water. RNA integrity was determined by running it on an agarose gel. Quantity was determined spectrophotometrically (Nanodrop).

### Methylated RNA immunoprecipitation (MeRIP)

The MeRIP procedure was modified from a protocol described in (Zeng et al., 2018). At least 50 μg of total RNA was randomly fragmented for 125 s at 90°C to approximately 200 nt using the NEBNext Magnesium RNA Fragmentation Module (New England Biolabs). Fragmented RNA was precipitated with 4 μl of GlycoBlue (Thermo Fisher), 20 μl of 3M sodium acetate pH 5.5 and 500 μl of 100% ethanol. Samples were freeze-thawed in dry ice three times and centrifuged at 12,000 x g and 4°C for 30 min. The pellet was washed with 1 ml of ice-cold 75% ethanol and centrifuged at 12,000 x g and 4°C for 10 min. RNA was resuspended in 20-100 μl of RNase-free water. Fragmentation was verified using agarose gel electrophoresis and RNA quantity was determined fluometrically using Qubit RNA high-sensitivity assay kit (Thermo Fisher).

At least 30 µg of fragmented RNA was used for each MeRIP. Fragmented RNA was incubated with 5 mg of anti-N6-methyladenosine antibody (Sigma-Aldrich, Cat. # MABE1006), 5 μl of RNase inhibitor (Lucigen) and 450 μl of IP buffer (150 mM NaCl, 10 mM Tris-HCl pH 7.5, 0.1% NP-40) at 4°C for 2 h with gentle rotation in a siliconized microtube (Millipore). Afterwards 25 µl of Protein A/G magnetic beads (Thermo Fisher) were added to each sample and incubated at 4°C for 2h. Samples were washed with washing buffer (500 mM NaCl, 10 mM Tris-HCl pH 7.5, 0.1% NP-40) and IP buffer twice. RNA was eluted by incubation of the beads with 200 μl of RLT buffer from the RNeasy Mini Kit (Qiagen) for 2 min. After bead separation using a magnetic rack, the eluate was transferred into a new reaction tube and 400 μl of 100% ethanol was added. The samples were transferred to an RNeasy MiniElute column and centrifuged at 12,000 rpm and room temperature. RNA was washed using 500 μl of RPE buffer from the RNeasy Mini Kit (Qiagen) and 500 μl of 80% ethanol. Columns were dried by centrifugation at 12,000 rpm and room temperature for 5 min. RNA was eluted with 20 μl of RNase-free water and yield was determined fluometrically using Qubit RNA high-sensitivity assay kit (Thermo Fisher).

### Sequencing library preparation and short-read sequencing

MeRIP sequencing (MeRIP-seq) libraries were prepared using the Ultra II directional RNA library preparation kit (NEB) as per manufacturer’s instructions for already fragmented samples. As input 5-50 ng RNA was used. Libraries were prepared for MeRIP input and elution samples. RNA-sequencing libraries were prepared from 500 ng total RNA using the Ultra II directional RNA library preparation kit (NEB) in combination with the NEBNext Poly(A) mRNA Magnetic Isolation Module (NEB). Single end sequencing of 76 bp was performed on NextSeq (Illumina).

### Data analysis

Trimmed sequencing reads were mapped to the *Arabidopsis* genome (TAIR10) using STAR ver. 2.7.5 (Dobin et al., 2013). m^6^A-sites on transcripts were identified via peak-calling by the R package exomePeak ver. 1.6.0. (Meng et al., 2013). Motif enrichment and metagene analyses were conducted using HOMER ver. 4.1 (Heinz et al., 2010). Methylated regions were annotated using the web-version of PAVIS (Huang et al., 2013). Cuffdiff ver. 2.1.1 was used to identify differentially expressed genes in the RNA-sequencing samples (Trapnell et al., 2012). Correlations between RNA-sequencing samples were determined using the R package corrplot ver. 0.84 (Wei and Simko, 2017). The DaPars algorithm (Xia et al., 2014) was employed to examine alternative polyadenylation. GO term enrichment analyses were performed using Panther ver. 16 (Thomas et al., 2003). The following R packages were used in this study: cowplot, extrafont, ggpubr, ggrepel, pheatmap, RColorBrewer, readr, readxl, reshape2, scales, tidyverse. Gene models were modified from images generated by Integrative genomics viewer (IGV) ver. 2.7.2 (Robinson et al., 2011).

### Motif analysis through deep learning approach

Enriched RNA motifs were identified using a deep learning approach described in (Koo and Ploenzke, 2021). In brief, positive class sequences were defined as combined m^6^A peaks in 3’UTR regions of transcripts in all samples with 200 nt up- and downstream extensions. Negative class sequences were defined as a randomly subsampled set of RNA-sequencing reads matched to the GC-content of positive class sequences. We randomly split the dataset into training, validation and test sets according to the fractions 0.7, 0.1 and 0.2, respectively. The total number of positive class sequences were 10163 and the total number of negative class sequences were 6196. A deep learning model, called ResidualBind (see Task 5 and 6 from Koo and Ploenzke (2021)), was employed to take one-hot encoded RNA sequences as input and predict the presence or absence of an m^6^A peak. The model was trained by minimizing the binary cross-entropy loss function with mini-batch stochastic gradient descent (100 sequences) for 100 epochs with Adam updates (Kingma and Ba, 2014) using default parameters. We set the learning rate to 0.001 and decayed it by a factor 0.2 when the AUROC on the validation set did not improve for five epochs. Saliency maps (Simonyan et al., 2013) were calculated by computing the gradients of the predictions with respect to the inputs and multiplied by the input sequence. Attribution maps were visualized as a sequence logo using Logomaker (Tareen and Kinney, 2019).

## SUPPLEMENTARY FIGURE LEGENDS

**Figure S1. Experimental conditions for identification of differential m^6^A RNA modification and validation**

**(A)** After 2 d of stratification at 4°C, indicated genotypes were incubated in respective light and temperature regimes to elicit developmental responses prior to tissue harvest. WT and *cry1cry2* mutant plants were grown in monochromatic blue light and WT and *phyB* mutant plants were grown in red light for 4 days prior to harvest. Etiolated WT plants were grown in darkness for 4 days and then subjected to a 24 h white light treatment to initiate de-etiolation. For shade and HT treatments, plants were grown in continuous white light for 4 days and treated with either shade or elevated temperature conditions for 24 h prior to harvest.

**(B)** Representative seedlings grown under the indicated conditions were imaged at the time point of harvest for RNA extraction.

**(C)** Hypocotyl length of the indicated genotypes in the indicated growth conditions. Asterisks indicate significance as determined by Student’s t-test. ****: *p* < 0.0001, ***: *p* < 0.0005, n= 17 – 21 (average: 19) seedlings were measured per condition.

**Figure S2. Quality control of RNA-seq dataset**

**(A)** Correlation plot of all RNA-seq replicates. Indicated correlation values represent Pearson’s rank correlation coefficient. Replicates were highly correlated (*r* ≥ 0.93).

**(B)** Differential expression of marker genes for respective conditions and the indicated genotypes. Expression values of significantly differentially expressed marker genes for sample pairs are shown as FPKM (fragment per kilo base million).

**Figure S3. m^6^A-methylation can be found in highly and lowly abundant transcripts** Significantly differentially expressed genes for all sample pairs are shown as FPKM. Colored dots indicate genes that were found to be uniquely methylated in either of the conditions. m^6^A was identified in significantly up- and downregulated genes.

**Figure S4. m^6^A is mainly found in the 5’UTR and around the TES**

Sequence logos of attribution maps generated by a deep learning model for a given mRNA sequence. Heights correspond to the importance of the nucleotide with a sign that corresponds to its impact on predictions. Analysis was performed on m^6^A-containing 3’UTR regions combined from all samples. RNA-seq reads were downsampled and GC-corrected to be within 95-105% of the GC-content of the 3’UTR regions and used as the null model. Identified UGUA motifs are shown for the top nine saliency maps (red boxes).

**Figure S5. m^6^A modification site and hypocotyl phenotypes of candidate genes**

**(A)** Gene models for candidate genes are represented by colored boxes. The identified site of m^6^A modification is marked with a dashed box.

**(B), (C)** Hypocotyl length of plants grown in shade (A) or HT (B) conditions are shown as bar plot. Error bars represent standard deviation. Red dots show the elongation response as calculated by the ratio of the average length in shade divided by the average length in ambient condition (N ≥ 12). Hypocotyl elongation of *lhy* and *ttl* mutant lines were similar to WT plants grown in shade. Hypocotyl length of *xpo1a* mutant lines were similar to WT plants when grown in HT, *bsk1* mutant lines exhibited stronger hypocotyl elongation than WT in HT.

**Figure S6. CRYs influence 3’UTR preference**

RNA-seq read counts of two exemplary genes, STO (A) and ALB3 (B) for WT B and *cry1cry2* mutant plants grown in blue light and the identified m^6^A-sites are shown together with a model of the 3’UTR.

