## Supplementary figures and images for "Light and temperature regulate m^6^A-RNA modification to regulate growth in plants"

### Supplemental Figure 1

#### Figure S1, Artz et al.

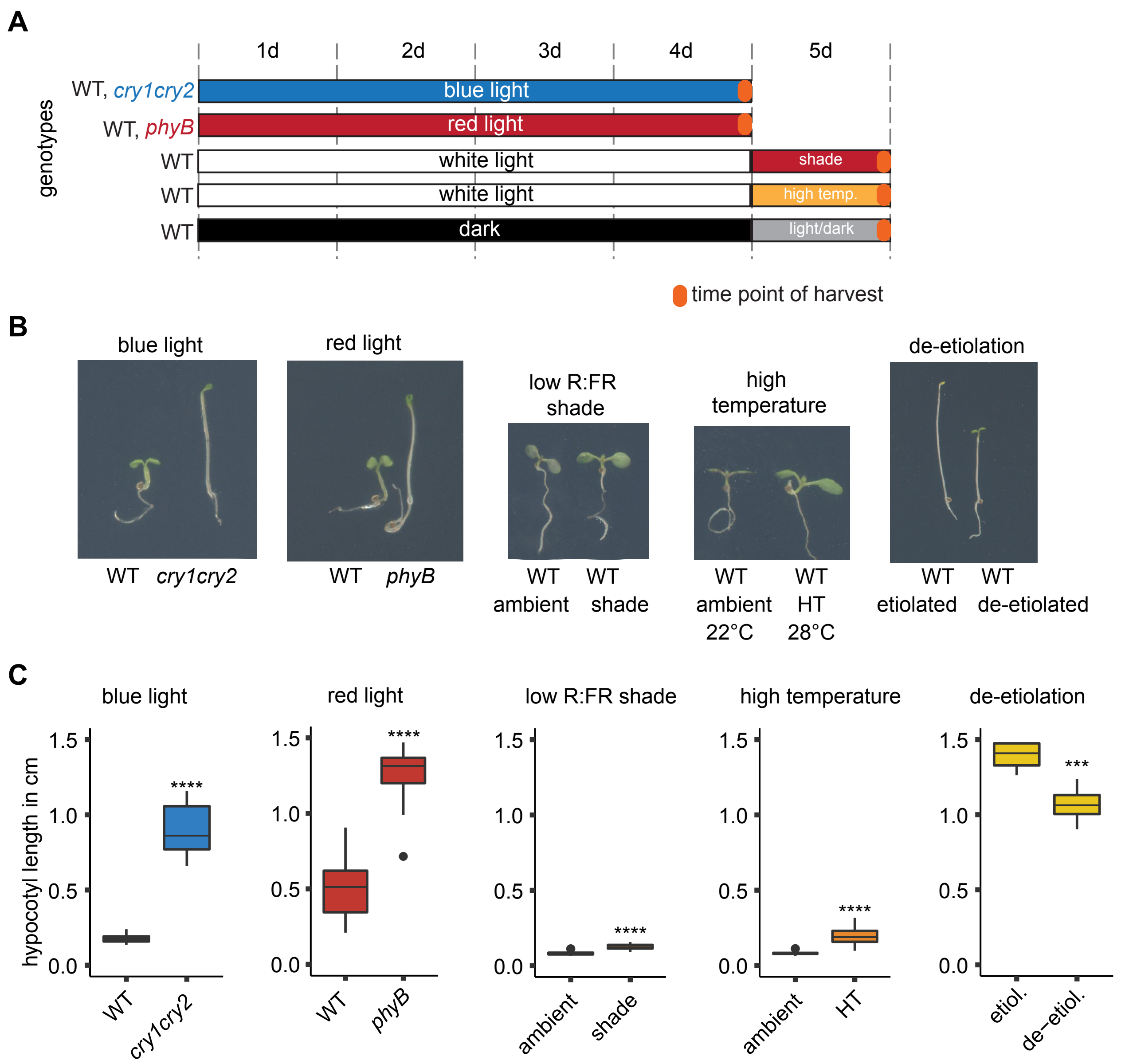

### Supplemental Figure 2

**Figure S2, Artz et al.**

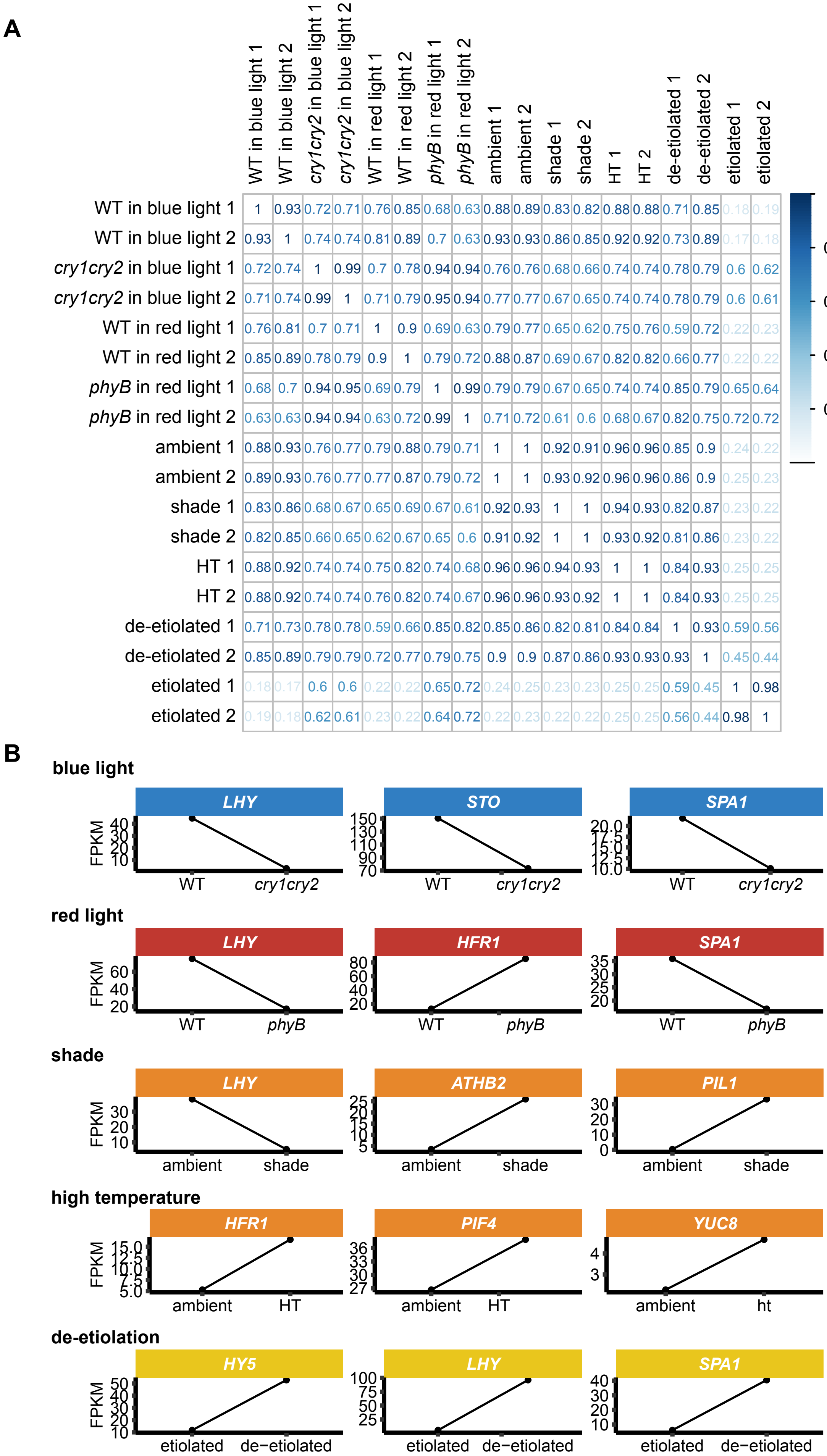

### Supplemental Figure 3

**Figure S3, Artz et al.**

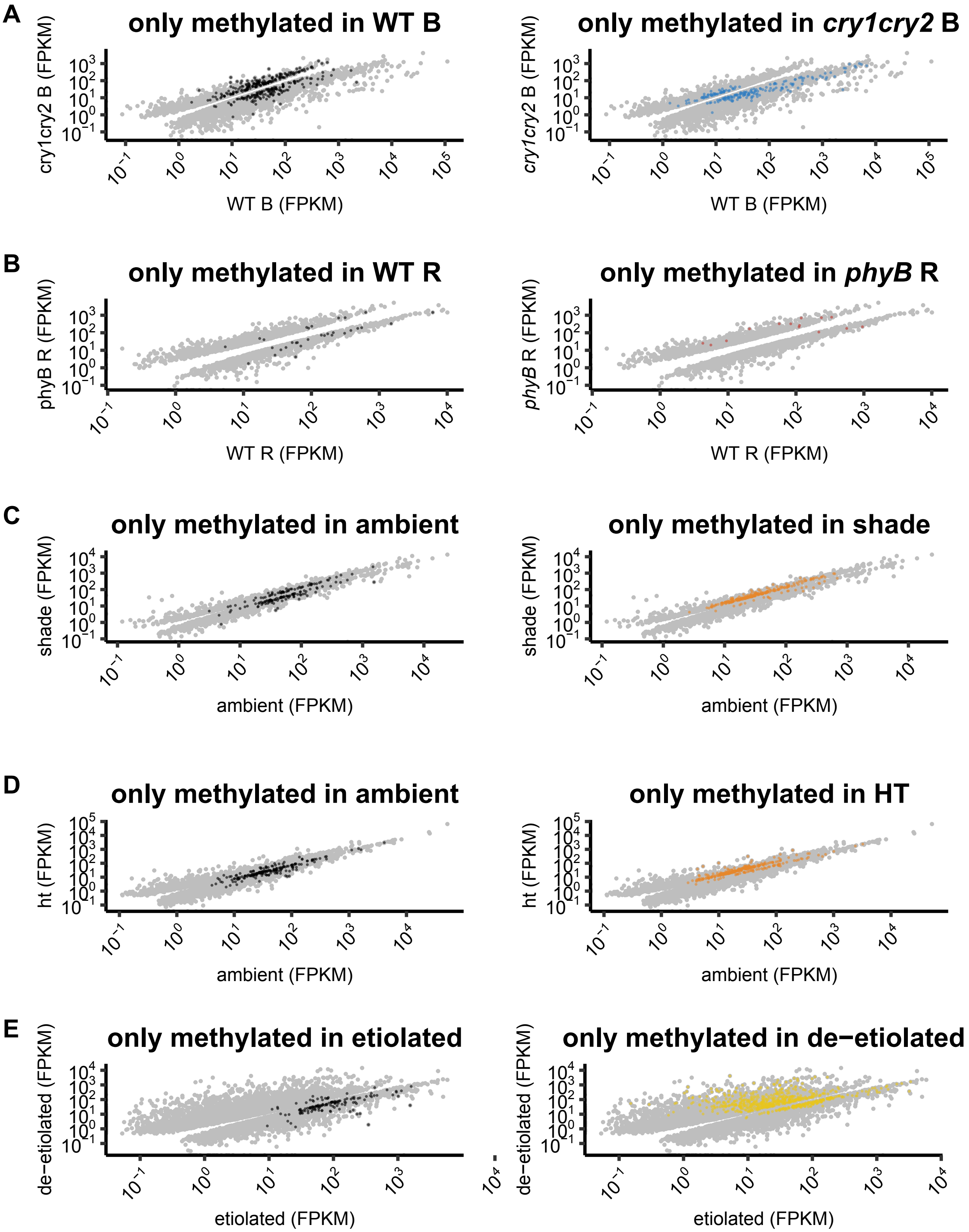

### Supplemental Figure 4

**Figure S4, Artz et al.**

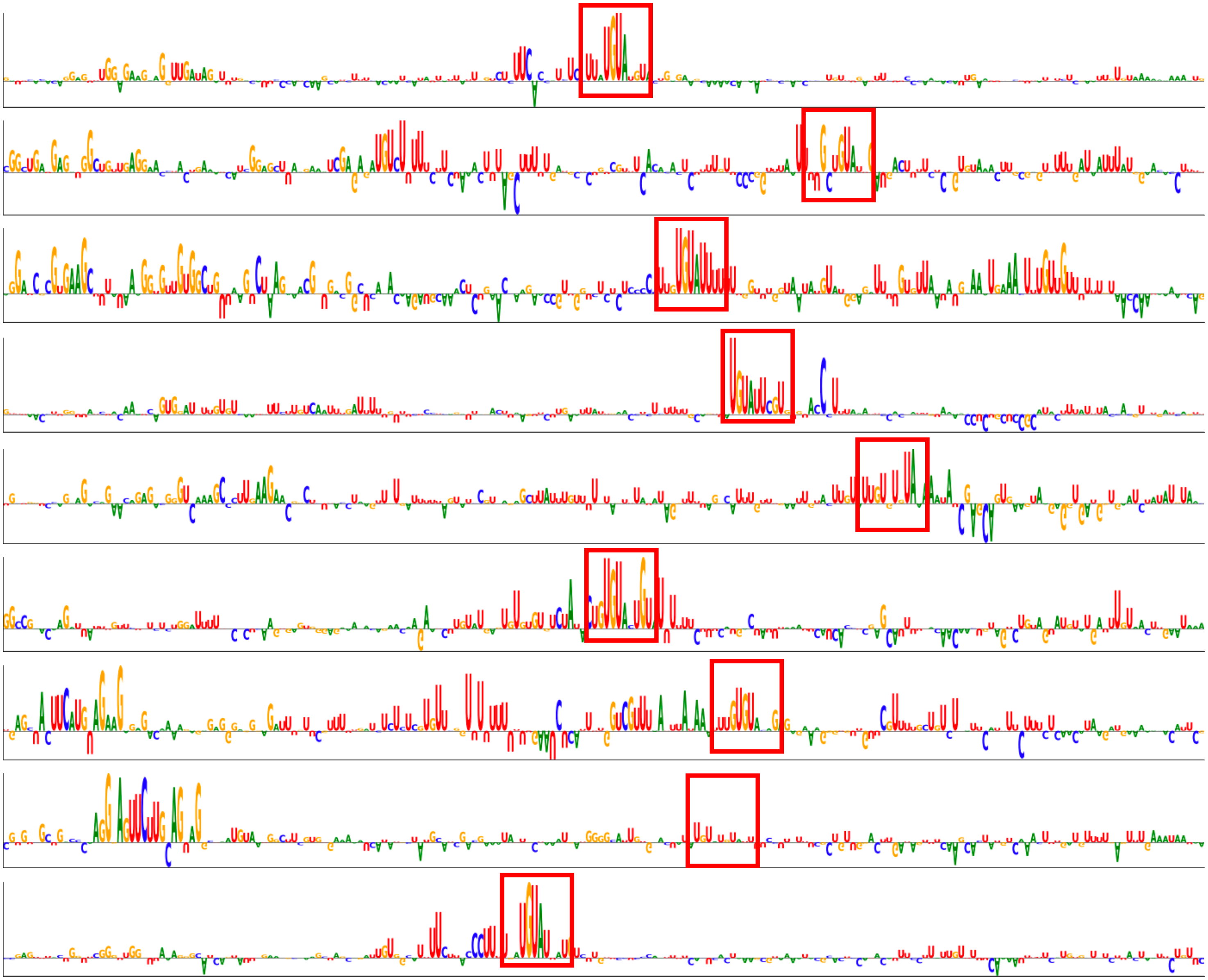

### Supplemental Figure 5

Figure S5, Artz et al.

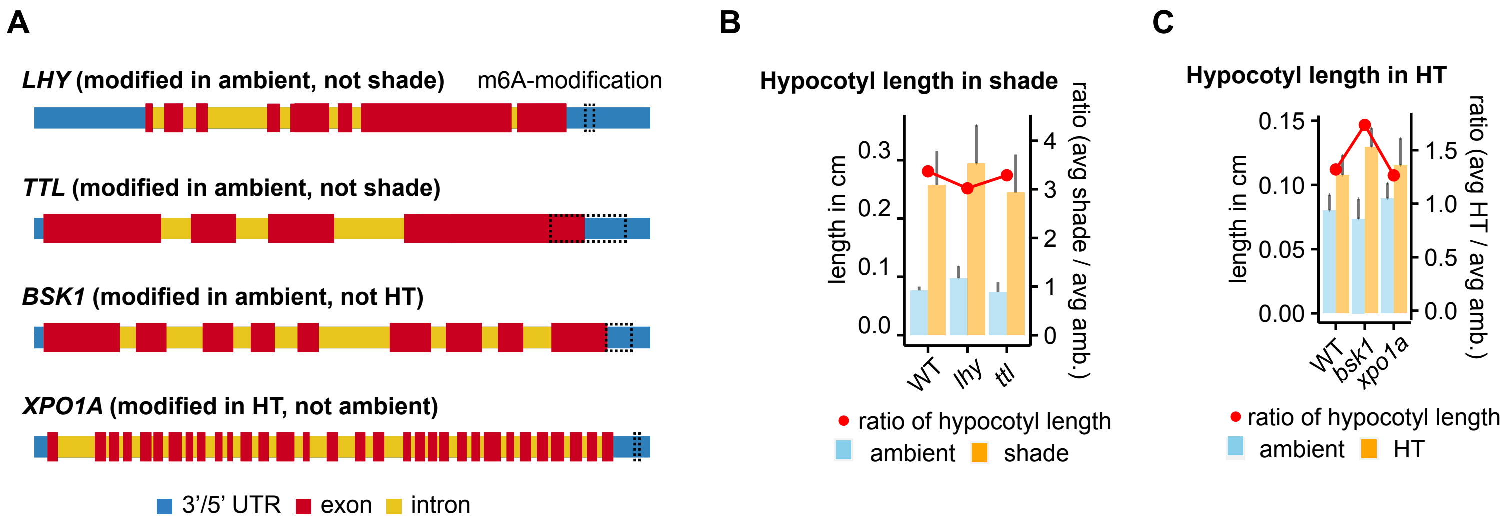

### Supplemental Figure 6

Figure S6, Artz et al.

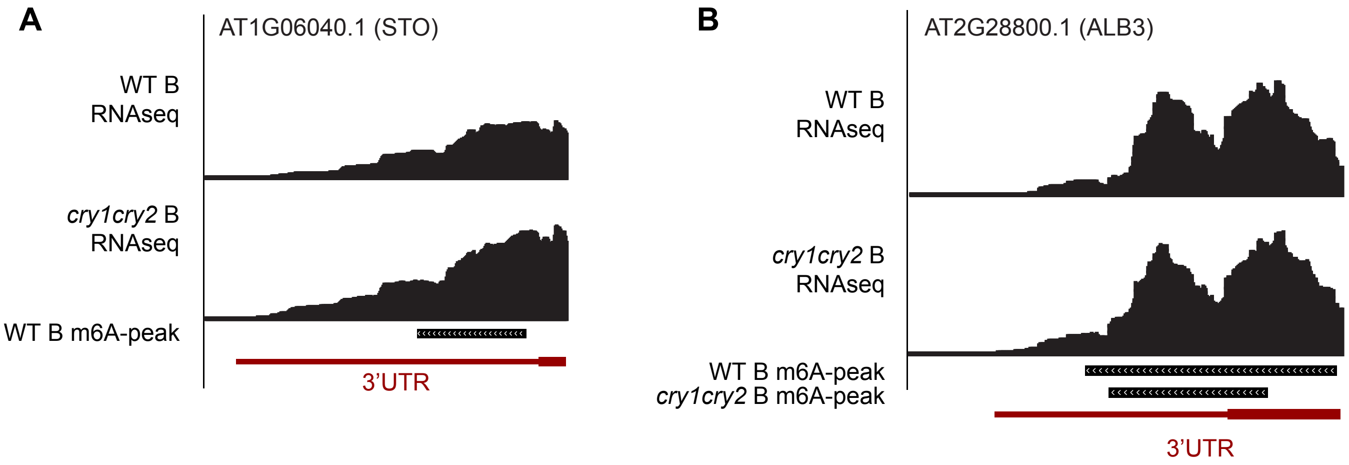
